# Cultivation-dependent effects of quorum sensing signals on a lactic acid and chain-elongating bacterium

**DOI:** 10.64898/2026.08.19.745728

**Authors:** Lena Depaz, Anouk Nys, Shanaya Scharloo, Celia Álvarez Fernández, Jana De Bodt, Josefien Van Landuyt, Jo De Vrieze, Ramon Ganigué

**Author notes:** Correspondence to: Ramon Ganigué, The University of Queensland, School of Chemical Engineering, G227+5F St Lucia, Brisbane, Australia; Webpage: https://chemeng.uq.edu.au/.

## Abstract

Microbial chain elongation enables the conversion of organic waste into higher-value products and is therefore a promising process for circular biomanufacturing. However, the microbial interactions governing chain elongation communities remain poorly understood. While quorum sensing has been extensively studied in the context of pathogens and model organisms, research on the perception of quorum-sensing molecules by non-model organisms and their effects within microbial consortia has remained limited. Here, *Lactiplantibacillus plantarum* and *Megasphaera elsdenii* were selected as representatives of two key functional guilds in chain elongation communities, namely lactic acid bacteria and chain-elongating bacteria. The effects of different exogenous quorum sensing molecules were evaluated in pure cultures and co-cultures using microtiter plates and serum bottles. Both organisms exhibited distinct molecule-dependent responses for both growth and biofilm formation. Moreover, the response of *M. elsdenii* was highly dependent on the supplied substrate. Despite changes in growth and/or biofilm formation, product yield and product spectra remained largely unaffected. Importantly, responses observed in pure cultures did not predict co-culture behavior, and no clear response to the tested molecules was detected in the co-culture grown in serum bottles. These findings demonstrate that responses to quorum sensing molecules are strongly dependent on the signal, substrate, microbial context, and cultivation conditions. These results highlight the limited predictive power of pure-culture assays for microbial communication in interacting communities and emphasize the importance of studying signal perception under process-relevant cultivation conditions.

**Key points:**

- Quorum sensing signals mainly affected growth and biofilm but not product spectra
- Quorum sensing effects were strongly substrate-dependent for *Megasphaera elsdenii*
- Cultivation conditions strongly determine quorum sensing effects, thus limiting applicability

**Graphical abstract:** 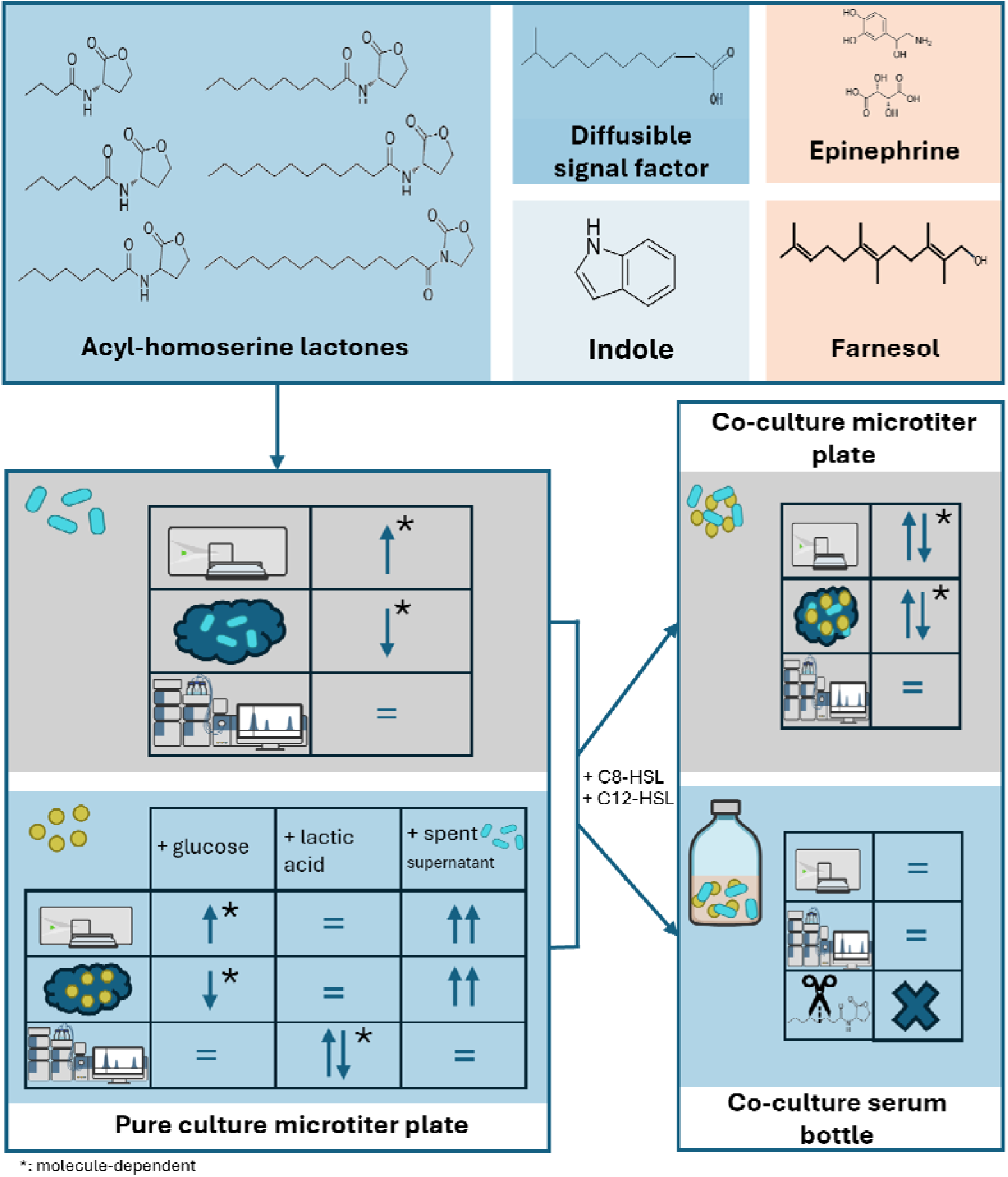

## INTRODUCTION

Anaerobic chain elongation has gained considerable interest in recent years as a promising biotechnological process to produce medium chain carboxylic acids (MCCA), from renewable resources using mixed microbial cultures (Roghair et al. 2016). These MCCA are attractive targets due to their broad range of applications, easier recovery compared to their shorter-chain counterparts (because of their higher hydrophobicity), and higher economic value compared to other precursor compounds (Agler et al. 2011). Chain-elongating bacteria (CEB) also produce biohydrogen as part of their metabolism, further enhancing the overall value of this process by generating multiple useful products (Angenent et al. 2016).

One of the key strengths of microbial chain elongation is the metabolic flexibility of most CEB, which allows them to utilize a wide variety of substrates (Candry and Ganigué 2021; Wang et al. 2026). Lactic acid bacteria (LAB) are frequently present in continuous chain-elongating communities and can be considered both a natural partner and a competitor in the process (Rombouts et al. 2020; Candry and Ganigué 2021; Ulčar et al. 2023). Substrate competition between LAB and CEB can occur for carbohydrates, but many CEB can also feed on the lactic acid (and/or acetate and ethanol) produced by the LAB, resulting in either a syntrophic interaction and/or a competitive interaction between the two groups (Ohnishi et al. 2022). The nature of this interaction is mostly attributed to the interplay of environmental conditions such as pH, temperature, abundance of each guild, substrate concentration and other environmental factors (Weiland-Bräuer 2021). There is a large body of evidence in nature that microbial communication plays a role in controlling tight ecological and trophic interactions, although little is known on how this may shape LAB-CEB interactions (Keller and Surette 2006)

It is known that many LAB utilize density-dependent signaling mechanisms, called quorum sensing, to coordinate gene expression for bacteriocin productions and other population-level behavior (Kuipers et al. 1998; Qian et al. 2023). However, it remains unclear whether CEB use similar quorum sensing mechanisms, and whether they can sense the signaling molecules produced by LAB and/or other members of typical chain-elongating communities. Classical quorum sensing systems include peptide-based signaling in many Gram-positive bacteria and acyl-homoserine lactone-based signaling in many Gram-negative bacteria (Fuqua and Greenberg 2002; Williams et al. 2007). Other signaling-related compounds, including autoinducer-2, diffusible signal factor, autoinducer-3-associated catecholamines, indole, and farnesol, have also been implicated in microbial interactions, stress responses, biofilm formation, virulence, and metabolic regulation (Li and Tian 2012; Deng et al. 2014; Kim and Park 2015; Liu et al. 2025a; Paul et al. 2025). One key interest here is biofilm formation, since biofilm systems provide protection against toxic or inhibiting substances and better biomass retention, which can protect the slow-growing chain-elongators during long term operation (De Groof et al. 2019; Candry et al. 2023). Importantly, microorganisms do not necessarily need to produce a given signaling molecule themselves to respond to it. Exogenous signaling compounds may act as ecological cues, stress signals, or modulators of interspecies interactions (Boyanova 2017; Venturi et al. 2018; Wei et al. 2026). Although quorum sensing-related compounds have been extensively studied in pathogenic bacteria, biofilms, and some anaerobic digestion systems (Vendeville et al. 2005; Anburajan et al. 2023; Liu et al. 2025b), their role in chain-elongating communities remains poorly understood. This knowledge gap is particularly relevant, due to the close metabolic interactions between chain-elongating community members and because of the known quorum sensing systems of LAB. Previous studies already gave an indication that the addition of quorum sensing molecules could lead to increased MCCA yield in chain-elongating communities, although mechanistical insight remains elusive (Li et al. 2021, 2023; Thau et al. 202).

In this study, the influence of exogenous quorum sensing molecules, specifically various AHLs, DSF, indole, epinephrine bitartate (as an autoinducer-3 analogue) and farnesol, were systematically investigated for their impact on the growth, biofilm formation and metabolite production of pure and co-cultures of *Lactiplantibacillus plantarum* and *Megasphaera elsdenii* as representatives of two major guilds in chain elongation communities, namely lactic acid bacteria and chain elongators. *L. plantarum* is a gram-positive, facultative homofermentative lactic acid bacterium that produces lactic acid, which can subsequently be used by the gram-negative *M. elsdenii,* a metabolically versatile organism capable of utilizing both carbohydrates and lactic acid. In the presence of both substrates, *M. elsdenii* appears to preferentially consume lactic acid over glucose, making this partnership particularly relevant for efficient MCCA and biohydrogen production (Weimer and Moen 2013; Ohnishi et al. 2022). By comparing organism-specific responses, substrate-dependent effects, and differences between simplified and more complex cultivation systems, we aimed to clarify whether quorum sensing-related compounds can influence the interaction between *L. plantarum* and *M. elsdenii* in the context of chain elongation.

## MATERIAL AND METHODS

### Strains and cultivation

*Lactiplantibacillus plantarum* (LMG6907) was obtained from the Belgian Coordinated Collections Of MicroOrganisms/LMG Bacteria (LMG, Belgium) and *Megasphaera elsdenii* (DSM 20460) was acquired from the Deutsche Sammlung von Mikroorganismen und Zellkulturen (DSMZ, Germany). The organisms were stored at -80°C in a 1:1 mixture of 80% glycerol and the following media (L^-1^): trypticase peptone 5g, yeast extract 0.4g, glucose.H_2_O 1.98g, K_2_HPO_4_ 2g, KH_2_PO_4_ 0.23g, CaCl_2_.2H_2_O 0.25g, NaCl 2.25g, MgCl_2_.6H_2_O 0.1g, NH_4_Cl 2.5g, Wolin modified mineral solution 10mL, resazurin solution (0.1% w/v) 1mL, selenite-tungstate solution 1mL, MES buffer 11.71g and vitamin K_1_ solution 0.2mL. The composition of the solutions can be found in supplementary information (section S1). The media was boiled, dispensed under N_2_/CO_2_ (90:10) and autoclaved. Wolin vitamin solution 10mL, Na_2_S.9H_2_O 0.30g, cysteine-HCl.H_2_O 0.30g and NaHCO_3_ 2g were added after autoclaving from sterile, anaerobic stocks with a N_2_ headspace. To obtain sterility, the stock solutions were filter-sterilized with 0.22 µm filters (Macherey-Nagel, Anderlecht, Belgium). After addition of all compounds, the media had a final pH of 6. The organisms were grown at 37°C without shaking and were transferred at least 2 times before the experiment. This media was also used for all subsequent experiments, except for *M. elsdenii* growing on lactic acid, where the glucose was substituted for 16.3mM DL-lactic acid, and *M. elsdenii* growing on the supernatant of *L. plantarum*, with no additional carbon source. However, after measuring the abiotic blanks after the experiment, it was observed that the actual lactic acid concentration was lower than intended (11.8mM instead of 16.3mM). This value is also used for the calculation of the element balances (section 2.7). For the experiment with spent supernatant, *L. plantarum* was grown for 24 hours with an inoculation ratio of 5% at 37°C on the media described above. After 24 hours, the culture was centrifuged at 5000g for 5 minutes and the supernatant was filter-sterilized using a 0.22 µm filter (Macherey-Nagel, Anderlecht, Belgium). The media was made anaerobic by letting it stand in an anaerobic chamber (N_2_/CO_2_, 90:10) (GP-Campus, Jacomex, TCPS NV, Rotselaar, Belgium) for 24 hours. The pH of the media with lactic acid and the spent supernatant was 5.6 and 5.7, respectively.

### Plate experiments

Plate experiments were performed in an anaerobic chamber (GP-Campus, Jacomex, TCPS NV, Rotselaar, Belgium) with an atmosphere of N_2_/CO_2_ (90:10) at a temperature of 37°C in untreated 96 well plates (Thermo Scientific, Fisher Scientific Belgium, Merelbeke-Melle, Belgium). Each organism was grown until the early stationary phase before the plate experiment, which was 24 hours after inoculation for *L. plantarum* and 48 hours for *M. elsdenii*.

### Pure cultures

Experiments with each organism were inoculated at an inoculation ratio of 5% in the media (section 2.1) with either no quorum sensing molecules (control), N-Butanoyl-L-homoserine lactone (C4-HSL, 10µM), N-hexanoyl-L-Homoserine lactone (C6-HSL, 10µM), N-octanoyl-L-Homoserine lactone (C8-HSL, 10µM), N-decanoyl-L-Homoserine lactone (C10-HSL, 10µM), N-dodecanoyl-L-Homoserine lactone (C12-HSL, 10µM), N-tetradecanoyl-L-Homoserine lactone (C14-HSL, 10µM), indole (500µM), epipephrine bitartate (10µM), farnesol (10µM) or cis-11-Methyl-2-dodecenoic acid (DSF, 1µM). The quorum sensing molecules C6-HSL, C8-HSL, C12-HSL, C14-HSL were acquired from Cayman Chemical (United States). The quorum sensing molecule C10-HSL was acquired from TargetMol (United States) and C4-HSL, indole, farnesol, DSF and epinephrine bitartate were acquired from Merck KGaA (Germany). The acylhomoserine lactones and DSF were stored as stock solutions in dimethyl sulfoxide (DMSO) at -20°C and diluted in phosphate buffered saline (PBS) as needed. The stock solutions were kept in small volumes in brown glass vials to ensure that the stock would not degrade, due to repeated thawing and freezing. Three plates per organism were made, which contained 5 replicates of each condition, for a total of 15 replicates per condition. To ensure that the DSMO had no effect on cell growth or biofilm formation, an additional plate for each organism was created with DMSO controls for DSF and each HSL, with 6 replicates per condition. The concentrations of the quorum sensing molecules and DMSO can be seen in the supplementary info (Table S1). The growth of the organisms, represented by the OD_620_ of all plates, was measured, and pictures were taken using the oCelloScope after 24 hours for *L. plantarum* and 48 hours for *M. elsdenii* (section 2.4.3). One plate was chosen for full well imaging. For each condition on each plate, the content of one well was stored at -20°C for measuring the metabolites (section 2.4.4), and the content of one well was fixated in ethanol and stored for flow cytometry (section 2.4.1). For each condition, there were, therefore, three replicates for measuring metabolites and three replicates for flow cytometry. The remaining wells were analyzed for biofilm formation using the crystal violet assay, as described in section 2.4.2, resulting in nine replicates per condition.

### Co-cultures

For the co-culture experiments, two quorum sensing molecules (C8-HSL and C12-HSL) were chosen for further investigation due to their prior usage in mixed-culture experiments and their response to either *L. plantarum* or *M. elsdenii* (Li et al. 2021; Li et al. 2023; Thau et al. 2025). Prior to inoculation, the cell count was measured using flow cytometry (section 2.4.1), and the organisms were diluted to ensure a 1:1 cell count inoculation. For each condition, four plates were made with 20 replicates per plate of the co-culture without added quorum sensing molecules (control), with added C8-HSL (10µM) and with added C12-HSL (10µM). Two of these plates were taken for analysis at 24 hours and the other two were taken for analysis at 48 hours, for a total of 40 replicates per condition per timepoint. Growth, metabolites, imaging and biofilm measurements were performed as for the pure culture experiments, except a total of four replicates were taken for measuring metabolites and four were taken for flow cytometry analysis.

#### Serum bottle experiments

Experiments in serum bottles were performed in triplicate in 200mL serum bottles with 100mL of medium as described in section 0 and an anaerobic headspace (90:10, N_2_/CO_2_). The inoculum was grown until early stationary phase and in the case of co-culturing, mixed to equal cell counts as measured with flow cytometry. A volume of 5mL of this mix was added to the bottles (5% inoculation ratio). For the pure cultures, they were mixed to equal cell counts with PBS. Two quorum sensing molecules (C8-HSL and C12-HSL) were chosen for further investigation, similar to the co-culture plate experiment (section 2.2.2). The bottles were incubated at 37°C. The experiments were sampled every 12 hours and terminated after 60 hours. First, the pressure was measured and a 3mL gas sample and a 2mL liquid sample were taken. A subsample was filter-sterilized (0.22 µm filter, Sartorius, Germany) and frozen for further analysis. Another subsample was taken for analysis of pH and OD_620_. The final subsample was stored in DNAse-free tubes at -20°C to ensure no contamination had taken place.

#### Analysis methods

##### Flow cytometry

The samples were diluted in PBS in 10-fold steps for a final volume of 200µL and stained with SYBR^®^ Green I (SG, 100x concentrate in 0.22 μm-filtered dimethyl sulfoxide, Invitrogen) for total cell counts. Staining was performed as described previously, with incubation for 20 minutes at 37°C in the dark (Goethals et al. 2025). Samples were analyzed immediately after incubation using an Attune™ Nxt flow cytometer (Fisher Scientific™, Germany) equipped with a violet (405 nm), blue (488 nm), yellow (561 nm) and red (637 nm) with Attune focusing fluid (Fisher Scientific™, Germany) as sheath fluid and an event rate between 200 and 2000 events s^-1^. The instrument performance was verified daily using Attune performance tracking beads (Fisher Scientific™, Germany).

##### Crystal violet assay

The crystal violet assay was performed as described previously with some modifications (Yang et al. 2014). Briefly, the planktonic cell density was measured (OD_620_) using a microtiter plate reader (Infinite M200, TECAN, Austria). The non-adherent cells were removed using tap water, and the biofilms were stained with 250 µL of a 0.1% w/v crystal violet solution. After 15 minutes of incubation at room temperature, the wells were rinsed 3 times. A volume of 250 µL of pure ethanol was added to each sample, and the absorbance was measured at OD_540_ using the same microtiter plate reader. The biofilm formation was reported as the crystal violet absorption at OD_540_ normalized to OD_620_ as a proxy for growth (Perry et al. 2025) (Eq. 1):

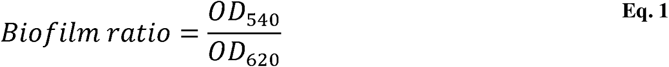

##### oCelloScope

Visualization of growth and cell attachment was done using the oCelloScope (Biosense, Denmark). The oCelloScope was in the anaerobic chamber to ensure no changes in cell morphology or density due to oxygen exposure. The oCelloscope UniExplorer (v. 11.0) was used to acquire 10 images/well after either 24 hours or after 48 hours depending on the experiment using the full well acquisition mode.

##### Analytical methods

Lactate, formate, acetate, propionate, butyrate and iso-buyrate were determined using ion chromatography. The ion chromatograph (930 Compact IC Flex, Metrohm, Switzerland) was equipped with a Metrosep organic acids 250/7.8 column, a Metrosep organic acids guard column/ 4.6 and an 850 IC conductivity detector (Metrohm, Switzerland). The anions were eluted with 1 mM H_2_SO_4_ at a flow rate of 0.5 mL⋅min^-1^. Glucose, valerate and hexanoate concentrations were determined by High-Performance Liquid Chromatography (HPLC). Samples were analyzed on a Shimadzu Prominence LC-2030C Plus with an RI detector (RID-20A). An isocratic method of 15 min on a Rezex ROA-Organic Acid H+ (8%) column (Phenomenex—part number 00F-0138-K0 + SecurityGuard Cartridge Kit (KJ0-4282) + SecurityGuard Cartridges Carbo-H 4 × 3.0 mm ID (AJ0-4490)) was used. The mobile phase was 5 mM sulfuric acid (Chem-lab CL00.2653.0050). The gaseous head space composition was analyzed using a Compact Gas Chromatograph (Global Analyser Solutions, Breda, Netherlands), equipped with a Molsieve 5A pre-column and Porabond column (CH_4_, O_2_, H_2_, and N_2_), and a Rt-Q-bond pre-column and column (CO_2_). Concentrations of gases were determined using a thermal conductivity detector. For the serum bottle experiments, the sample at timepoint 60 hours was additionally analysed for C3 to C8 carboxylic acids (including isoforms C4 to C6) by gas chromatography (GC) with flame ionization detection (FID) as described earlier (Candry et al. 2020a).

#### Quorum sensing inhibition assay

The quorum sensing inhibition assay was performed as described previously (Darai and Pelyuntha 2025). *Chromobacterium violaceum* (ATCC 12472) was obtained from American Type Culture Collection (ATCC, United States of America). *C. violaceum* was grown in LB at 28°C with shaking. Briefly, 5mL of LB inoculated with 100µL *C. violaceum* ATCC 12472 was treated with 200 µL of cell-free supernatant or fresh LB (negative control) and incubated at 28°C for 24 hours. After incubation, the culture broth was centrifuged to separate the bacterial cells and remove the supernatant. The resulting pellet was resuspended in 1 mL DMSO and vortexed vigorously to extract violacein. The absorbance of the violacein-DMSO mixture was measured at 585 nm using a plate reader. All experiments were performed in triplicate. The percentage of violaceum inhibition was calculated as follows:

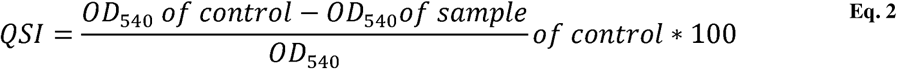

#### Statistical analysis

For each response variable, descriptive statistics were calculated for each condition, including the number of replicates, mean, standard deviation, standard error of the mean, median, minimum, and maximum. Prior to selecting the primary inferential test, a one-way linear model was fitted for each dataset using condition as a categorical explanatory variable. Model diagnostics were evaluated using residual Q-Q plots, residuals-versus-fitted plots, Shapiro–Wilk tests on model residuals, and the Brown–Forsythe test for heterogeneity of variance. The Brown–Forsythe test was implemented as Levene’s test using deviations from the group medians.

For conditions where sample size was low, formal assessment of normality and variance homogeneity was considered unreliable and statistical power was limited and therefore a Steel-type rank-based permutation test with max-statistic adjustment was used. These results were interpreted cautiously, with emphasis on effect direction and magnitude, rather than statistical significance alone. For conditions with sufficient sample size, parametric treatment-versus-control comparisons were used when residual normality was acceptable. Welch’s two-sided independent t-tests versus the untreated control with Holm correction were used as the default parametric treatment-versus-control approach for both equal variance and unequal variance because the same reporting strategy was preferred across organisms and datasets. If the residual normality was not acceptable, Steel-type rank-based permutation test with max-statistic adjustment was used. Statistical analysis was performed using the SciPy package (version 1.16.3) in Python (version 3.13.9). Detailed information on the statistical analysis results can be found in the supplementary information per experiment.

#### Calculations

Estimations of soluble EPS, bound EPS and biomass were done based on a chemical oxygen demand (COD) balance. A sample of 20 mL was taken and centrifuged at 4°C at 10956g for 20 minutes. The supernatant was taken and stored at -20°C until further analysis (soluble EPS+media EPS). The pellet was resuspended in 20mL milliQ water and heated at 60°C for 30 minutes to loosen any bound EPS from cells. The heated tubes were then centrifuged at 4°C at 10956rpm for 20 minutes. The supernatant was then stored at -20°C for further analysis (bound EPS). Finally, the remaining pellet was resuspended in 20mL of milliQ and stored at -20°C until further analysis (biomass). The COD of the soluble EPS, bound EPS and biomass were measured using nanocolor kits (Macherey-Nagel, Belgium). The COD balance was made as follows:

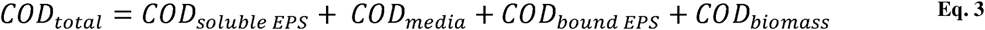

For the electron balances, molar product concentrations were converted to electron equivalents per liter by multiplying the concentration with the amount of electron liberated during the oxidation of the component (Candry et al. 2020b). The following values were utilized: glucose (24 mol e^-^/mol), lactate (12 mol e^-^/mol), formate (3 mol e^-^/mol), acetate (8 mol e^-^/mol), propionate (14 mol e^-^/mol), isobutyrate (20 mol e^-^/mol), butyrate (20 mol e^-^/mol), valerate (26 mol e^-^/mol), isovalerate (26 mol e^-^/mol), caproate (32 mol e^-^/mol) and ethanol (12 mol e^-^/mol). Only glucose or lactic acid was considered as consumed. Biomass (CH_1.8_O_0.5_N_0.2_, 4.2 mol e-/mol) and hydrogen gas (2 mol e-/ mol) were not considered in the balance unless specifically mentioned. For the experiments in plates, only glucose, lactate, formate, acetate, propionate, butyrate and isobutyrate could reliably be measured due to the high dilution rate of the samples. Therefore, an estimate of the missing products was made. For this, biomass of *M. elsdenii* was assumed to be 10% of the consumed carbon when growing on glucose and 3% when growing on lactate (see section S1.4) and the remaining unknown products were assumed to be valerate, caproate, CO_2_ and H_2_ based on known yields and product spectra for growth on glucose and on lactic acid (Weimer and Moen 2013). The estimation was made by simultaneously solving elemental balances (carbon, hydrogen, oxygen and electron balances), calculated as a linear least-squares problem with only non-negative solutions (Eq. 4) using the SciPy package (version 1.16.3) in Python (version 3.13.9).

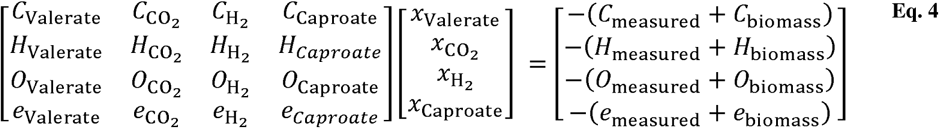

## RESULTS

### Exogenous quorum sensing molecule addition to L. plantarum

*L. plantarum* was grown for 24 hours with various quorum sensing molecules to assess the impact of the molecule on its growth, adhesion to a micro-titer plate and its metabolite production. For growth, the OD_620_ values ranged between 0.266 ± 0.024 (control) and 0.278 ± 0.016 (C12-HSL). The biofilm ratio (OD_540_/OD_620_) values ranged between 0.73 ± 0.186 (epinephrine bitartate) and 0.973 ± 0.254 (control). In general, the biofilm ratio had a higher variance than the measured growth. All quorum sensing molecules, except C8-HSL, C10-HSL and indole, resulted in significantly improved growth (Figure 1, Table S6.). This resulted in a slightly reduced biofilm ratio in all conditions, although this decrease was not statistically significant. C12-HSL and C14-HSL had the highest increase for growth: 23% (0.278 ± 0.016) and 22% increase (0.276 ± 0.010) respectively compared to the control (0.226 ± 0.024). On the other hand, epinephrine bitartate and farnesol had the greatest decrease in biofilm formation with a 25% decrease (0.730 ± 0.186) and a 24% decrease (0.742 ± 0.168) respectively compared to the control (0.973 ± 0.254). The condition with the least amount of change was C8-HSL, with no difference in either biofilm ratio (0.941 ± 0.178) or growth (0.238 ± 0.030). Changes in response were not considered to be a result of the DMSO concentration (Figure S4).

**Figure 1.**
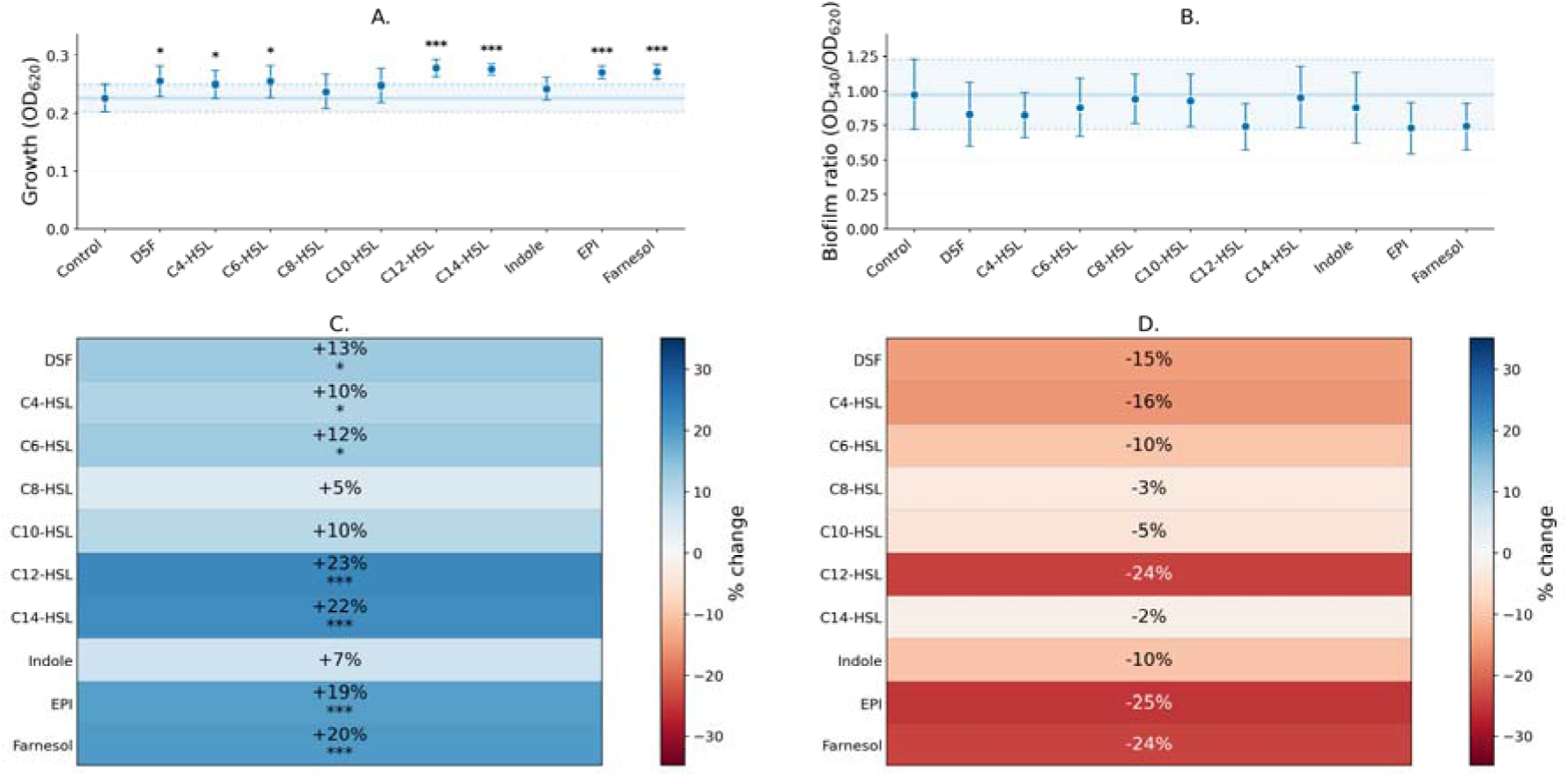
Growth (A, n=15) and biofilm ratio (B, n=9) of *L. plantarum* after 24 hours with different quorum sensing molecules added or without any quorum sensing molecules added (control). Error bars indicate standard deviation. The solid line indicates the average value of the condition without exogenous quorum sensing molecules (control), while the dotted lines and the blue band indicate the mean ± 1 standard deviation. Growth compared to the control (C) and biofilm ratio compared to the control (D) show the change in percent compared to the control. * indicates significant changes compared to the untreated control (p < 0.05), ** indicates p < 0.01 and *** indicates p < 0.001.

Lactic acid was the sole metabolic product and none of the exogenous quorum sensing molecules had a significant impact on lactic acid production, with concentrations ranging from 14.5mM to 16.4mM (Figure S2). No acetate or ethanol was detected for any of the samples. The electron balance for tested compounds and the control closed between 72% (farnesol) and 82% (control), not taking biomass into account (Figure S3).

### Exogenous quorum sensing molecule addition to M. elsdenii

The different quorum sensing molecules caused substrate-specific differences for *M. elsdenii* as seen in both growth and normalized biofilm formation (Figure 2). Like *L. plantarum*, biofilm ratio had a higher variance than growth. *M. elsdenii* growing on the spent supernatant of *L. plantarum* led to the lowest biofilm ratio (control: 1.165 ± 0.187), followed by *M. elsdenii* growing on glucose (control: 3.489 ± 1.600) (Figure S7). *M. elsdenii* growing on lactic acid had the highest biofilm ratio of all conditions (control: 3.841 ± 0.861). The highest growth was observed with glucose (control: 0.324 ± 0.083), followed by spent supernatant (control: 0.241 ± 0.027). The lactic acid condition had the least growth (control: 0.168 ± 0.051).

**Figure 2.**
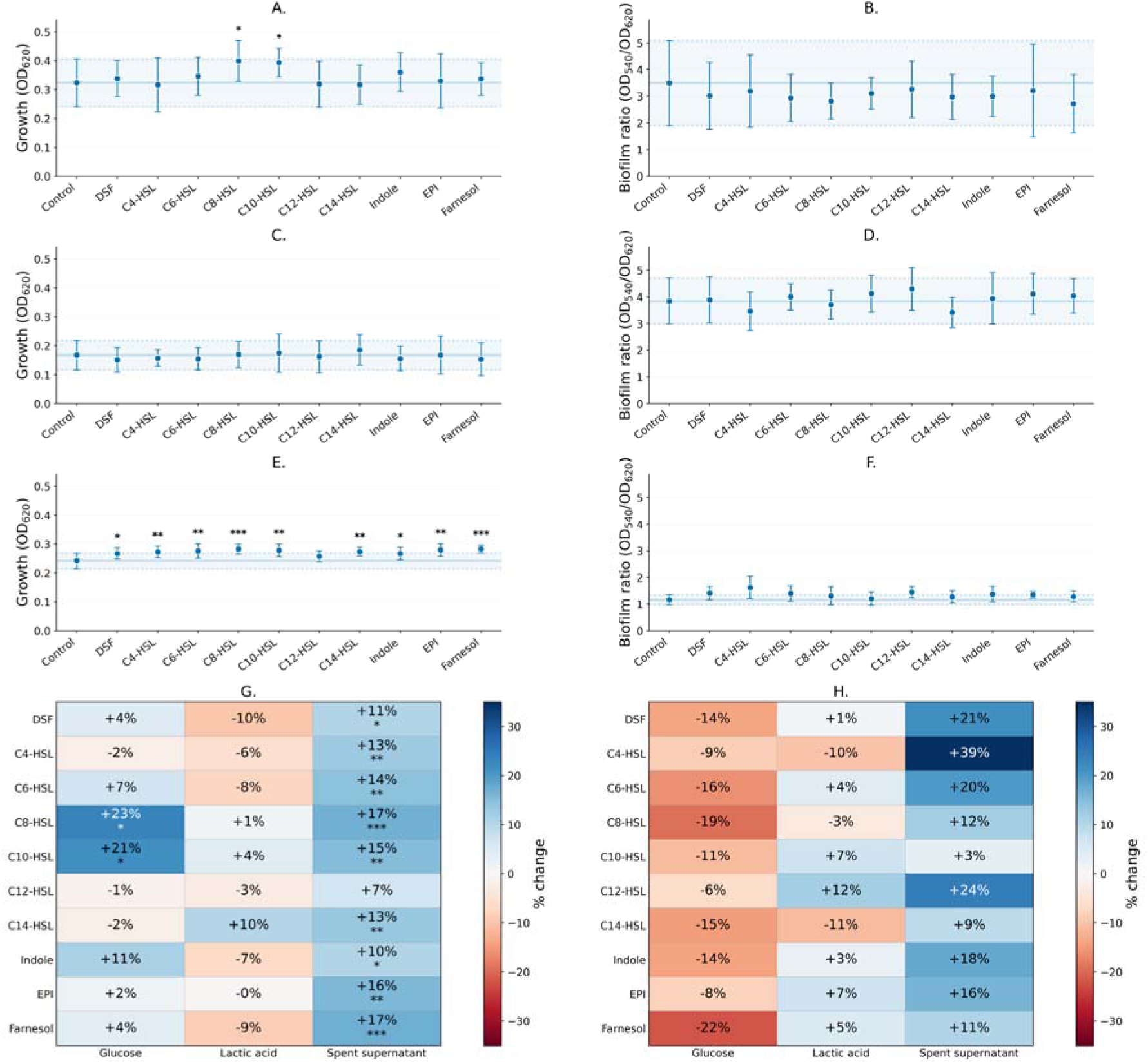
Growth of *M. elsdenii* after 48 hours with different quorum sensing molecules added or without any quorum sensing molecules added (control) on glucose (A), lactic acid (C) and spent supernatant (E). Biofilm of *M. elsdenii* after 48 hours with different quorum sensing molecules added or without any quorum sensing molecules added (control) on glucose (B), lactic acid (D) and spent supernatant (F). Error bars indicate standard deviation. The solid line indicates the average value of the condition without exogenous quorum sensing molecules (control), while the dotted lines and the blue band indicate the mean ± 1 standard deviation. Growth compared to the control (C) and biofilm ratio compared to the control (D) show the change in percent compared to the control. * indicates significant changes compared to the untreated control (p < 0.05), ** indicates p < 0.01 and *** indicates p < 0.001.

The addition of C8-HSL (0.399 ± 0.071) and C10-HSL (0.393 ± 0.049) resulted in a significant increase (23% and 21% respectively) in growth of *M. elsdenii* using glucose as substrate, while the growth on lactic acid was not significantly impacted by any of the quorum sensing molecules. This is in stark contrast to the growth on the spent supernatant. This could be partly due to the higher concentration of lactate in the spent supernatant. In the spent medium conditions, all the exogenous quorum sensing molecules, except for C12-HSL, led to a statistically significant increase in growth as indicated by OD_620_ (Figure 2). As for the biofilm ratio, none of the exogenous quorum sensing molecules led to any significant changes when growing *M. elsdenii* on glucose or lactic acid. Although no significant changes were observed, the addition of any quorum sensing molecule to *M. elsdenii* growing on glucose led to a general slight decrease in its biofilm production with a decrease ranging from 6% to 22% compared to the control, as also seen for *L. plantarum*. When growing on lactic acid, the trend of increased growth at the expense of biofilm formation is true for most quorum sensing molecules, except for C4-HSL (growth decreased 6% and biofilm decreased 10%) and C10-HSL (growth increased 4% and biofilm increased 7%). No significant changes in biofilm ratio were observed when growing on the spent supernatant, but all quorum sensing molecules led to an increase in biofilm ratio (+3% to +37%). Changes in response were not considered to be a result of the DMSO concentration (Figure S11).

When growing on glucose, the metabolite profile consisted on a molar basis mainly of acetate (49 – 62.5%) and butyrate (26.3 – 33.6%), with small amounts of isobutyrate (3 – 4.2%) and formate (8.4 –13.2%) (Figure B, Figure S8A). While growing on lactic acid (Figure D, Figure S8B), the profile shifted to acetate (32.4 – 39.2%), propionate (21.4 – 30.2%) and butyrate (21.9 – 30.1%) as the main products, with an increased fraction of isobutyrate (8.9 –13.2%) and a slightly decreased amount of formate (3.3 - 6.0%) compared to the conditions with glucose. A similar profile is seen when growing on the spent supernatant (Figure F, figure S8C), with acetate (31.6 – 33.9%), propionate (19.5% - 22.5%) and butyrate (32.6 – 37.4%) as the main products and small amounts of isobutyrate (5.1 – 5.7%) and formate (5.5 – 5.8%).

There was no significant difference in metabolite profiles between the controls and the experiments with addition of exogenous quorum sensing molecules, neither on glucose (Figure 3A and B) nor on lactic acid (Figure 3C and D). When growing on spent supernatant, the addition of the quorum sensing molecules caused in most conditions an increase in acetate (between -1% and +11%) and propionate (between +3% and +18%) at the expense of butyrate (between -5% and -17%) and isobutyrate (between -2% and -18%), except for C12-HSL. However, there was only a statistically significant decrease in butyrate for C10-HSL (3.13 ± 0.29mM) compared to the control (3.79 ± 0.13mM). The metabolite profile for C12-HSL was almost identical to the untreated control, which is in line with the fact that only a slight increase in growth was observed (+7% versus control). The conditions with longer-chain AHLs showed a more pronounced decrease in butyrate (from -10% to -17%) compared to the shorter chain AHLs (from -5% to -9%). The condition with lactate resulted in lower overall production compared to the spent supernatant, which is likely due to the lower than intended initial concentration of lactate (section 2.1).

**Figure 3.**
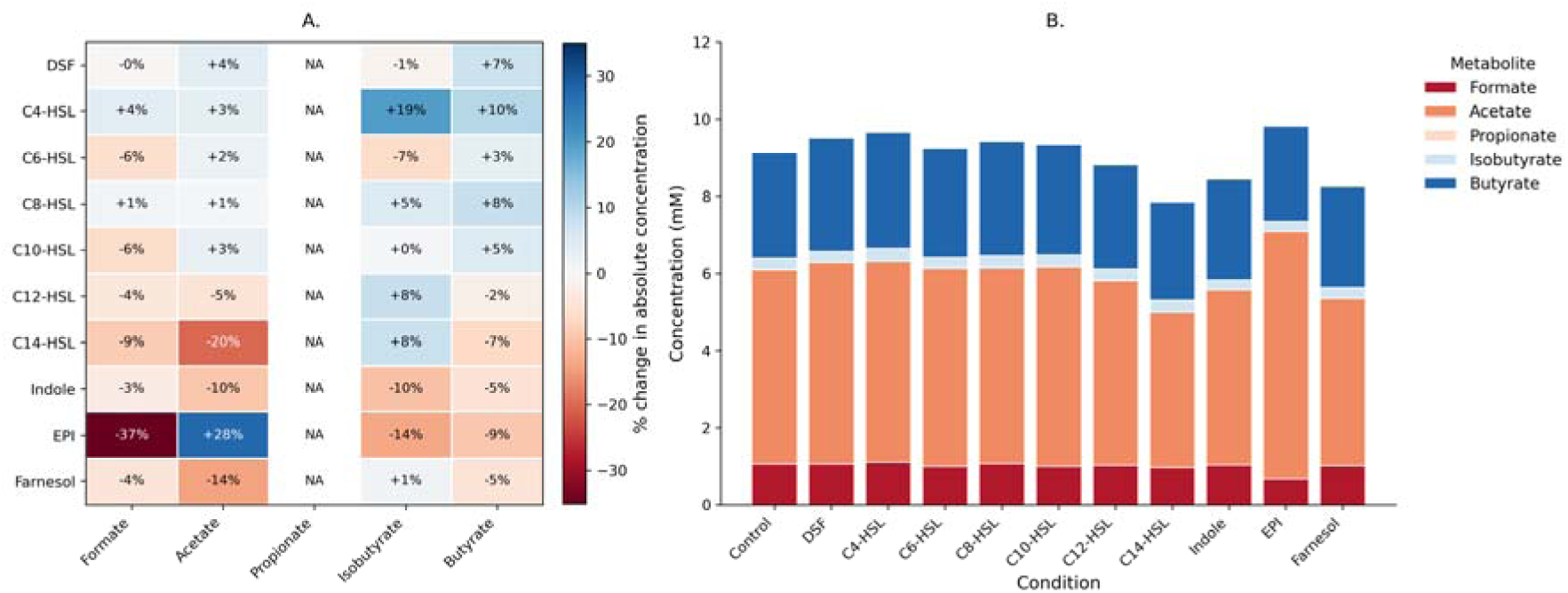
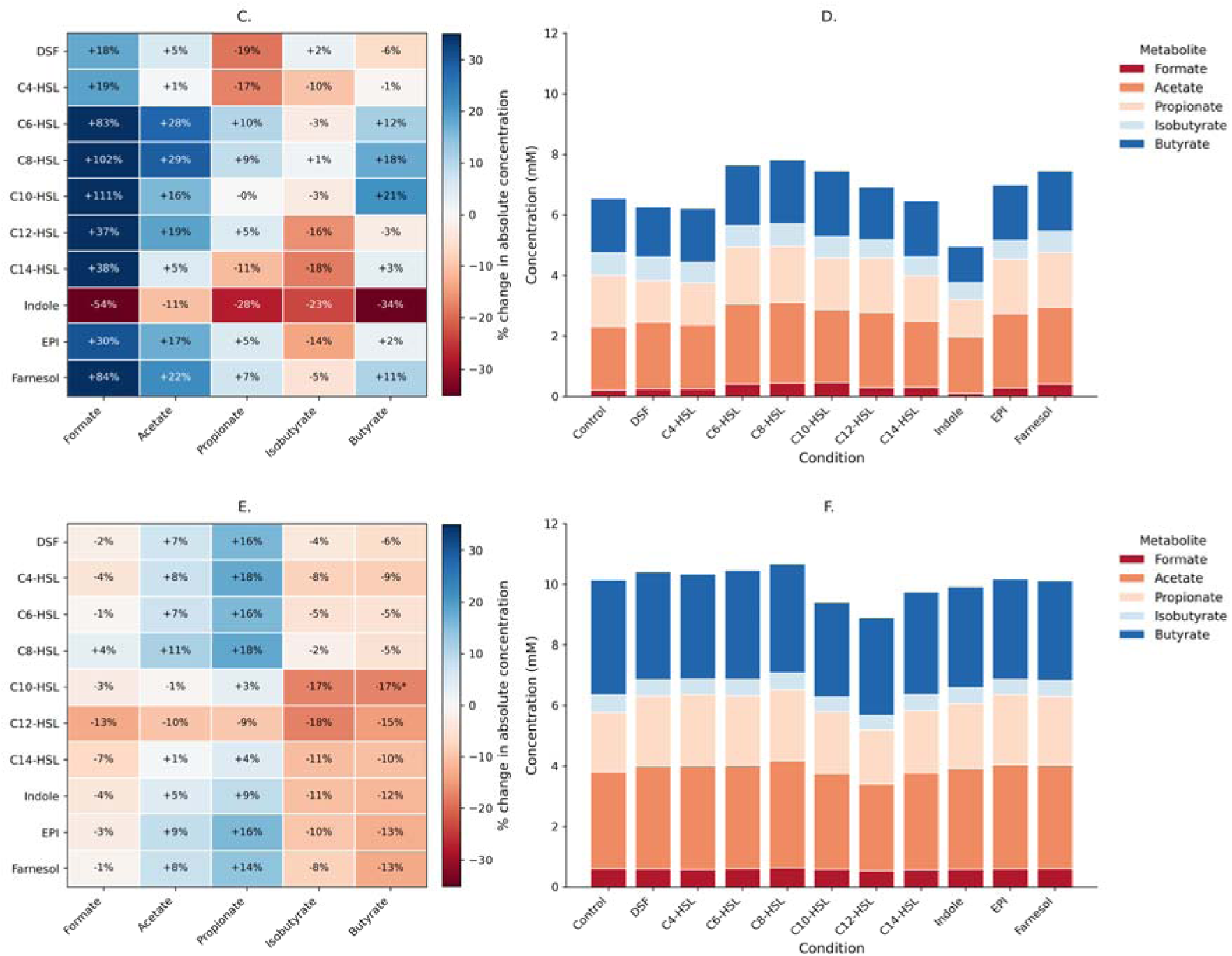
Metabolite production of *M. elsdenii* after 48 hours. Change in absolute concentration of metabolites with quorum sensing molecules compared to the control without exogenous quorum sensing molecules when growing on glucose (A), lactic acid (C) and the spent supernatant of *L. plantarum* (E). Absolute metabolite production with or without exogenous quorum sensing molecules when growing on glucose (B), lactic acid (D) and the spent supernatant of *L. plantarum* (F). * indicates significant changes compared to the untreated control (p < 0.05)

When considering all measured products, glucose (Figure S9A) showed a low level of electron balance closure (40% to 48%), whereas the electron balance for the spent supernatant (Figure S9C) closed the best with a range of 63% (C12-HSL) to 75% (C8-HSL). This is without considering the produced hydrogen and biomass. The condition with lactic acid (Figure S9B) displays a similar level of closure compared to the condition with the spent supernatant with a range of 61% to 75%, except for indole, which only closed for 48% when growing on lactic acid. Based on the calculated values for the missing products for glucose, around 2.6 ± 0.1mM of biomass, 1.8 ± 0.01mM of valerate, 1.4 ± 0.01mM of caproate, 17 ± 0.04mM of CO_2_ and 16.5 ± 0.4mM of H_2_ would be needed to close the element balances (Figure S10A, Table S14). The missing products for growth on lactate would be 0.32 ± 0.05mM of biomass, 0.57 ± 0.20mM of valerate, 0.45 ± 0.14mM of caproate, 9.94 _±_ 0.51mM of CO_2_ and 9.80 ± 0.56mM of H_2_ based on the element balances (Figure S10B, Table S15). Finally, the missing products for *M. elsdenii* growing on the spent supernatant of *L. plantarum* would be 0.39 ± 0.03mM of biomass, 0.59 _±_ 0.12mM of valerate, 0.47 ± 0.1mM of caproate, 13.27 ± 0.28mM of CO_2_ and 12.98 ± 0.29mM of H_2_ (Figure S10C, Table S16). In all conditions, a 10 times dilution resulted in concentration of valerate and hexanoate below the quantifiable limit of the HPLC, which is 0.49mM for valerate and 0.43mM for hexanoate.

### Exogenous QS molecule addition to the co-culture

The response of the microorganisms to quorum sensing additions in a co-culture were further explored for two selected molecules, C8-HSL and C12-HSL. An initial screening on 96-well plates showed that at 24 hours, the addition of C8-HSL resulted in a statistically significant decrease in growth, while the addition of C12-HSL resulted in a statistically significant increase in biofilm ratio (Figure 4A). In this experiment, the condition with exogenous C8-HSL showed a decrease in growth and an increase in biofilm compared to the control also after 48 hours, but the condition with exogenous C12-HSL had a lower biofilm ratio and significantly more growth, despite the significantly positive effect on the biofilm formation of *M. elsdenii* growing on the *L. plantarum* supernatant observed before (Figure 2).

**Figure 4.**
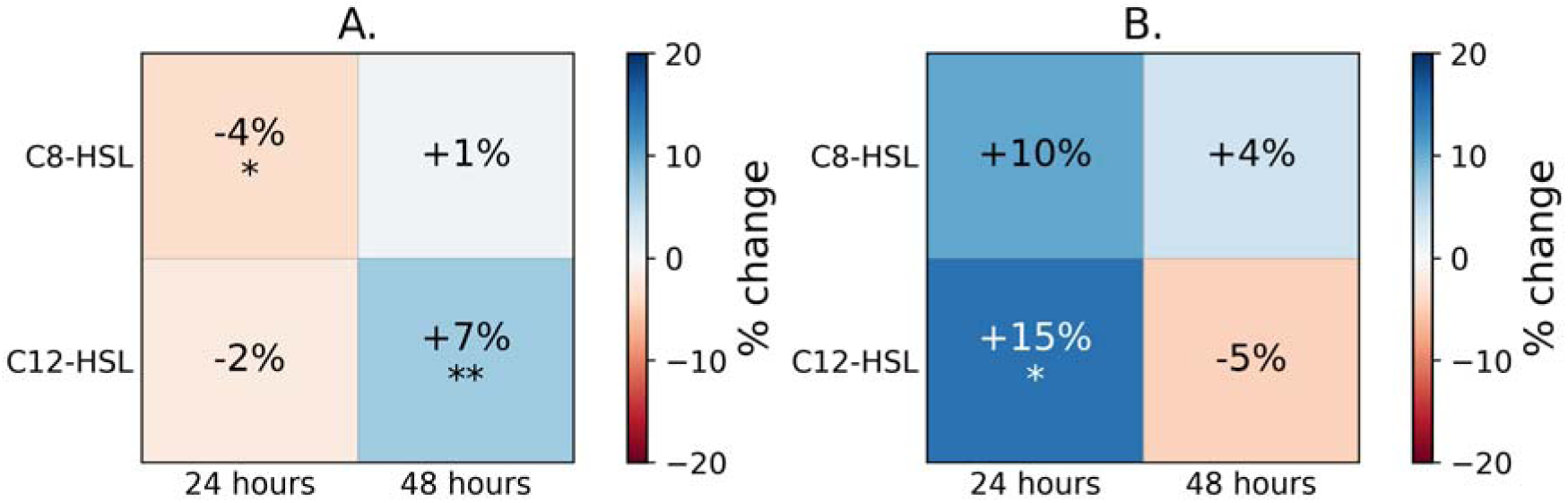
Change in growth (A) and biofilm ratio (B) of the co-culture of *L. plantarum* and *M. elsdenii* growing with C8-HSL and C12-HSL compared to the control without exogenous quorum sensing molecules after 24 hours and 48 hours. * indicates significant changes compared to the untreated control (p < 0.05), ** indicates p < 0.01.

Acetate, propionate and butyrate were all produced concurrently at 24 hours, indicating both glucose and lactic acid consumption (Figure 5). Acetate (5.3 ± 0.3 mM for the control, 5.2 ± 0.26 mM for the condition with C8-HSL and 5.3 ± 0.47 mM for the condition with C12-HSL) and propionate (2.5 ± 0.21 mM for the control, 2.31 ± 0.11 mM for the condition with C8-HSL and 2.33 ± 0.32 mM for the condition with C12-HSL) peaked at 24 hours. Between 24 hours and 48 hours, acetate and propionate concentrations decreased, while the concentration of butyrate increased monotonically. Propionate was no longer detected at 48 hours, while acetate decreased to 0.61 ± 0.56mM for the control, 1.38 ± 0.09mM for C8-HSL and 1.38 ± 0.09mM for C12-HSL. Neither lactic acid nor glucose were observed after 24 hours. Isobutyrate was only observed at 48 hours, at a concentration of 0.44 ± 0.05mM for the control, 0.46 ± 0.03mM for C8-HSL and 0.47 ± 0.05mM for C12-HSL. At 24 hours, the electron balances close for 52% for the control and the condition with exogenous C12-HSL and closes for 50% for the condition with exogenous C8-HSL (Figure S14). This level of closure decreased to 41%, 44% and 44% for the control, exogenous C8-HSL and exogenous C12-HSL respectively at 48 hours, indicating an increase in not-accounted for products.

**Figure 5.**
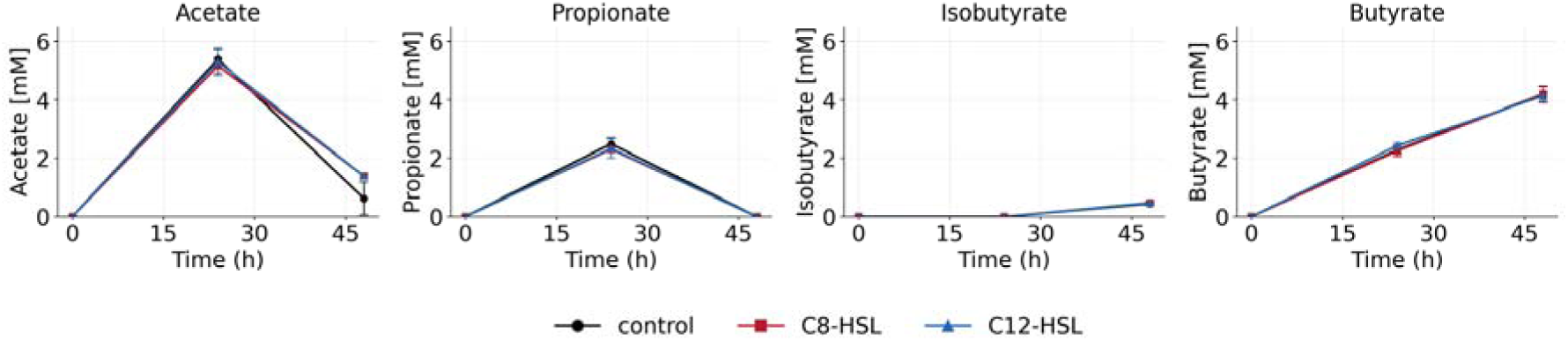
Concentration of acetate, propionate, isobutyrate and butyrate in the co-culture over time without added quorum sensing molecules 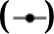, with added C8-HSL 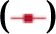 and with added C12-HSL 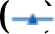 * indicates significant values (p < 0.05)

To gain deeper insights, a batch experiment was conducted in serum bottles to increase the resolution and follow the evolution of growth and metabolites (Figure 6). For the condition without added quorum sensing molecules, glucose was completely depleted after 36 hours, although no measurement is available at 24h making it difficult to pinpoint the exact moment when the substrate was depleted. Low levels of lactate were measured at 12 (0.2 ± 0.09mM) and 24 hours (0.48 ± 0.46mM), after which it was no longer detected. Acetate (1.07 ± 0.06mM) and propionate (0.04 ± 0.07mM) both peaked at 24 hours like the plate experiments, but both were observed at lower levels compared to the plates. Butyrate also peaked at 24 hours (3.25 ± 0.09mM), and in this case the concentrations were in line with the plate experiments. Isobutyrate (0.32 ± 0.02 mM) was detected at 24h, earlier than the 48 hours in the plates. Caproate (0.71 ± 0.71mM) was detected at 12 hours and remained stable at 36 hours and 48 hours. Valerate was detected using the HPLC but below the quantifiable limit. Since the stated products, including H_2_, only allowed the electron balance to close for 70%, the sample at 60 hours was additionally measured using VFA-GC. This showed isovalerate (2.39 ± 0.10mM) as an additional product. This closed the electron balance to 91% for the control, 100% with C8-HSL and 95% with C12-HSL (Figure S15).

**Figure 6.**
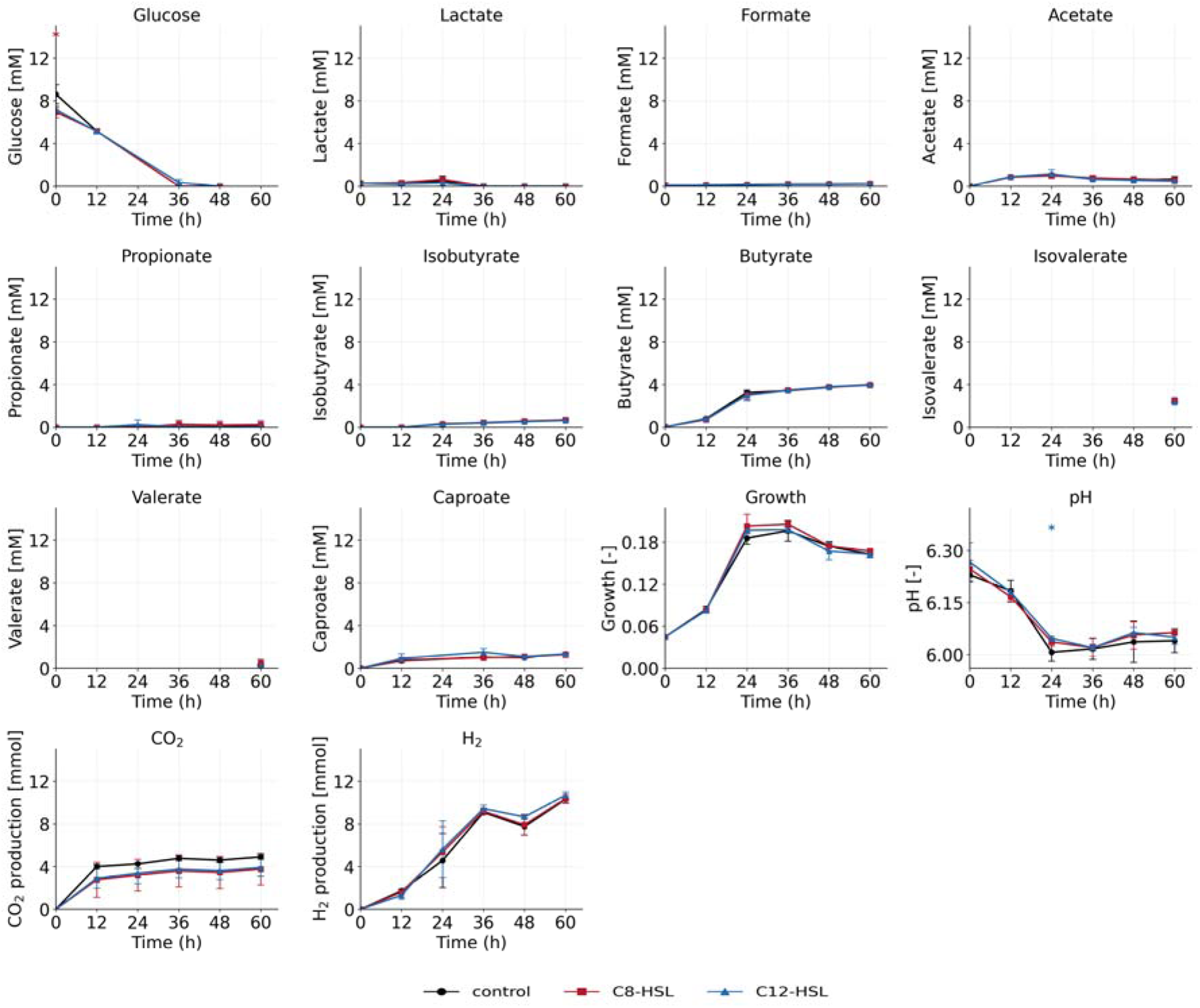
Concentration of glucose, lactate, acetate, propionate, isobutyrate, butyrate, isovalerate, valerate, caproate, growth, CO_2_ and H_2_ in the serum bottles of the co-culture over time without added quorum sensing molecules 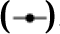, with added C8-HSL 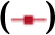 and with added C12-HSL 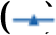. * indicates significant values (p < 0.05)

Neither C8-HSL nor C12-HSL addition led to a significant difference in metabolic profile or growth profile. As opposed to what was seen in the microtiter plate experiments, propionate did not decrease with C8-HSL, C12-HSL or without quorum sensing molecules.

There was observable microbial adherence to the glass of the serum bottle in all conditions when growing in co-culture, which was repeatedly disturbed by sampling. An estimate of biofilm formation after 60 hours was made based on EPS and biomass concentrations (Table S19). For the co-culture, C8-HSL and C12-HSL, no significant changes in either soluble EPS, bound EPS or biomass were observed. Therefore, it is assumed that there is no significant difference in biofilm formation.

The bio-detector *Chromobacterium violaceum* was used to analyze the quenching potential of the supernatant of the pure cultures of *L. plantarum*, *M. elsdenii* and the co-culture, which had a quorum sensing inhibition of 17%, 9% and 2% respectively (Table S20). This indicates that the individual pure cultures have a greater capacity to degrade or interfere with AHLs than the combination of the organisms.

## DISCUSSION

### Exogenous signaling molecules affect pure cultures in an organism- and substrate-dependent manner

Quorum sensing is increasingly understood as a broader and more complex phenomenon than the classical Gram-positive versus Gram-negative framework suggests. Although true quorum sensing remains difficult to prove experimentally, current research indicates that a broader range of organisms can at least detect quorum-sensing molecules, even if they do not produce them themselves (Rajput and Kumar 2017; Venturi et al. 2018; Schwieters and Ahmer 2025). Many reported effects may therefore reflect signal perception, perhaps as an environmental stressor, rather than signal production and associated quorum sensing response.

In these experiments, the response of *L. plantarum* to the added quorum sensing molecules was characterized primarily by increased OD_620_. Because lactate concentrations were not significantly affected, the increased OD_620_ does not appear to reflect a major rerouting of the carbon metabolism toward alternative fermentation products. Instead, the response may indicate improved biomass yield, altered maintenance energy requirements, increased stress tolerance, or changes in cell morphology or aggregation (Buck et al. 2009; Li et al. 2025). The simultaneous decrease in normalized crystal violet staining for selected compounds, including C12-HSL, epinephrine, and farnesol, further suggests that these molecules may alter surface properties, rather than simply increasing growth. This is consistent with the idea that some exogenous signaling molecules are perceived by *L. plantarum* as environmental cues or stressors, potentially affecting membrane composition, hydrophobicity, or extracellular polymer production (Wang et al. 2017; Haddaji et al. 2017). The native quorum sensing system of *L. plantarum* has been shown to be quite sensitive to environmental parameters (including temperature, pH and nutrient level, among others) and the presence of other microorganisms (Liu et al. 2017; Man and Xiang 2023; Liu et al. 2026). Therefore, the observed responses could be similarly dependent on the specific environmental parameters tested.

*M. elsdenii* displayed a strong substrate-dependent response. The limited effects observed during growth on glucose or lactate suggest that exogenous signaling molecules alone did not generally stimulate *M. elsdenii* growth under the tested conditions. However, when *M. elsdenii* was grown on spent *L. plantarum* supernatant, almost all tested molecules except C12-HSL increased OD_620_. This indicates that the components present in the spent supernatant of *L. plantarum* may have strongly altered the response of *M. elsdenii* to exogenous signaling-related compounds, although the exact compounds or mechanisms responsible are unknown. *L. plantarum* could have depleted certain nutrients in the condition with spent supernatant, which have been shown in *Pseudomonas synxantha* to lower the AHL activation threshold (Alcalde et al. 2026), or introduced other stressors like bacteriocins, that could likewise lower the activation threshold. Spent supernatant contains not only residual carbon sources, but also organic acids, peptides, vitamins, extracellular polymers, and other metabolites produced by *L. plantarum*. These components could alter membrane physiology, redox balance, nutrient limitation, stress sensitivity or induce or suppress the native quorum sensing system, thereby changing the apparent effect of the added compounds (Noike et al. 2002; Almeida et al. 2016; Tan et al. 2024). Other stressors, like bacteriocins, could have especially played a role in the co-cultures, since the presence of other organisms can trigger bacteriocin production (Liu et al. 2017; Man and Xiang 2023; Liu et al. 2026). Additionally, the concentration of lactate was higher in the spent supernatant compared to the condition with lactate, which could have acted as a stressor and lowered the activation threshold. *M. elsdenii* did not react to all AHLs to the same degree for neither normalized biofilm formation nor growth, indicating that sensitivity depends not only on the presence of the quorum sensing molecule, but also on its specific structural properties, including chain length. The strongest effect on growth was observed with C8-HSL and C10-HSL, while the strongest effect on biofilm ratio was observed with C4-HSL and C12-HSL. Earlier studies with C8-HSL addition have also noted an increase in abundance in *Megasphaera* spp. and other chain-elongating organisms like *Clostridia*, although this could have been an indirect effect (Zhao et al. 2024).

When looking at the metabolite profile of *M. elsdenii* cultures grown on spent supernatant, proportionally less butyrate was observed compared to acetate and propionate. Acetate production from lactate yields more ATP (1mol ATP/mol lactate) than butyrate (0.5mol ATP/mol lactate) and an increase in acetate production is often linked to an increase in NADH generation (Prabhu et al. 2012). This is partially in line with the observed growth profiles: for most conditions, the increase in acetate and propionate was concomitant with higher biomass, except for C10-HSL and C14-HSL, where only small changes in acetate concentration were observed despite large changes in OD_620_. This could be partly attributed to unaccounted fermentation products, in line with the low degree of closure of the electron balances (Figure S9). On the spent supernatant, the highest concentration of valerate and caproate is calculated for C10-HSL, C12-HSL and C14-HSL, which is in line with the biggest decrease in butyrate compared to the control. However, C10-HSL and C14-HSL led to an increase in growth, while growth was unaffected by the addition of C12-HSL. The calculated missing products of *M. elsdenii* growing on lactate and spent supernatant result in very similar product spectra profiles, but lower gas production was expected with spent supernatant (10mM H_2_) compared to lactate (13mM H_2_). Moreover, less hydrogen production is predicted compared to the condition with glucose (17mM), which is in line with other observations (Ohnishi et al. 2022). As stated in section 3.2, around 2.6 _±_ 0.1mM of biomass, 1.8 _±_ 0.01mM of valerate, 1.4 _±_ 0.01mM of caproate, 17 _±_ 0.04mM of CO_2_ and 16.5 _±_ 0.4mM of H_2_ would be needed to close the element balances in the plates with *M. elsdenii* growing on glucose (Figure S10A, Table S14). This is in line with the measured values in the serum bottles with *M. elsdenii* growing on glucose, except for the presence of isovalerate instead of valerate and higher predicted values for CO_2_ compared to the measurements (Figure S12). Isovalerate and isobutyrate can be produced when *M. elsdenii* performs amino acid fermentation, likely due to glucose and/or lactic acid depletion in combination with the high amount of yeast extract and peptone available in the media (Wallace 1986). However, this is difficult to verify. This also implies that the actual degree of closure is lower than calculated, as yeast extract and peptone consumption were not considered.

### Increasing complexity reduces the apparent effect of the quorum sensing molecules

Two levels of complexity were added from the micro-titer, pure culture experiments: co-culture cultivation in micro-titer plates followed by co-culture cultivation in serum bottles. The addition of C8-HSL and C12-HSL affected one or both micro-organisms in pure culture, yet these effects were not observed in co-cultures, neither in microtiter plates nor serum bottle incubations. In pure culture, the addition of C8-HSL had no effect on either growth or biofilm ratio of *L. plantarum* (Figure 1) but resulted in significant higher growth of *M. elsdenii* growing on glucose and spent supernatant (Figure 2). The addition of C12-HSL resulted in significantly higher growth of *L. plantarum* (Figure 1), but no impact on the growth of *M. elsdenii* (Figure 2). It was therefore expected that in the condition with C8-HSL, the product spectra would resemble the condition with growth on glucose, while the condition with C12-HSL would have resulted in a higher peak of lactate before its consumption. However, there was no difference in growth or metabolites when comparing the control with the addition of C8-HSL or C12-HSL. One possible explanation is that the experimental design could have changed the apparent response. Microtiter plates have a high surface-area-to-volume ratio, and hydrophobic signaling molecules, such as longer-chain AHLs, may partition into or adsorb onto plastic surfaces, potentially altering their effective concentration over time (Wang et al. 2025). Differences in material properties may have affected cell attachment, biofilm formation, and signal availability. The repeated sampling may have also interfered with the cell attachment, thereby masking a potential effect of the quorum sensing molecules. Beyond the type of material and their surface properties, serum bottle incubations differ from microtiter plates in volume, headspace composition, mixing regime, sampling procedure and geometry of the vessel, which could all impact the outcome of cultivations in subtle, but potentially important ways (Candry et al. 2018). However, this would not explain the difference in response of the co-culture compared to the pure cultures when grown in the same microtiter plates.

Another, non-mutually exclusive explanation is that the effect of the added signaling molecules was altered or masked by the higher biological complexity of the co-culture system, as also seen in the pure cultures of *M. elsdenii* growing on glucose or lactic acid compared to the spent supernatant. In the co-culture, the added molecules are embedded in a dynamic system involving lactate exchange, competition for residual carbohydrates, pH changes, product inhibition, and potential shifts in thermodynamic feasibility for example (Noike et al. 2002; Candry et al. 2018; Procházková et al. 2024). These changes in conditions may have masked the direct impact of the quorum sensing molecule by stronger metabolic and thermodynamic constraints.

The lack of a clear response to C8-HSL and C12-HSL in the co-culture, despite clear differences in the pure culture, has important ramifications for understanding the role of quorum sensing in more complex chain elongation cultures, and more broadly in any complex biological system. A major limitation of the co-culture experiments is that total growth and metabolite production could not reveal whether the added molecules affected *L. plantarum*, *M. elsdenii*, or both, since biomass quantification parameters like OD are a bulk aggregate of all species. Species-specific qPCR or time-resolved metaproteomics would link changes in community composition to metabolite profiles. These insights could allow to better understand given responses to quorum sensing molecules, or their lack thereof. Beyond controlled systems with exogenous addition, these tools could also allow better understanding of QS in more complex mixed culture systems.

### Quorum quenching did not appear to mediate the observed interaction under the tested conditions

Various *Lactobacilli*, including various strains of *L. plantarum*, can interfere with AHL production, likely through pH-related effects and the production of lactate or other volatile fatty acids like acetate effect (Kiymaci et al. 2018; Rana et al. 2020). Therefore, quorum sensing inhibition was measured at the end of the experiment for the pure cultures and the co-culture. The observed quorum sensing inhibition of 17% in this experiment (Table S20) for *L. plantarum* is in line with the expectations, given previous reported quenching of lactic acid bacteria ranging from 0 to 20%, and *L. plantarum* specifically being shown as a potential quorum quenching micro-organism (Lv et al. 2021; Darai and Pelyuntha 2025). On the other hand, the co-culture displayed very low to non-existent quorum sensing inhibition (2%), even compared to the pure culture of *M. elsdenii* (9%). Both the low quorum sensing inhibition of the co-culture and the 9% quorum sensing inhibition of the pure culture of *M. elsdenii* were unexpected. The low inhibition in the co-culture is evidence that quorum-quenching activity is context dependent and cannot simply be predicted from the properties of the individual species. The lack of quorum quenching in the co-culture despite quenching occurring in the pure culture of *L. plantarum* could indicate that the metabolic interaction between the two organisms counteracted the observed quenching activity of *L. plantarum*. Specifically, lactate was only observed at low concentrations (0.20 ± 0.07mM at its peak at 12 hours) which could indicate reduced production by *L. plantarum* or rapid consumption by *M. elsdenii*, thereby preventing the quorum-quenching associated accumulation of lactate (Darai and Pelyuntha 2025).

## CONCLUSIONS

Exogenous quorum-sensing-related molecules can affect the growth and surface-associated behavior of *L. plantarum* and *M. elsdenii*, but these effects are strongly dependent on organism, substrate, and cultivation format. In pure cultures, *L. plantarum* responded to several AHLs, including C12-HSL and C14-HSL, and other signaling-related compounds with increases in growth (+5% to +23%) and decreases in biofilm ratio (−5% to -25%), whereas lactate production remained largely unchanged. *M. elsdenii* showed a substrate-dependent response with limited effects when consuming glucose or lactate alone, but a broader stimulation when grown on spent *L. plantarum* supernatant, resulting in increased growth (+7% to +17%) and increased biofilm ratio (+3% to +39%). Responses observed in microtiter plates did not consistently translate to co-cultures in serum bottles, although it could not be verified whether this was due to biological reasons or the variations in experimental conditions. Overall, exogenous quorum-sensing-related compounds can modulate micro-organism phenotype, but pure-culture assays can have limited predictive power for microbial communication in interacting communities. Studying signal perception under process-relevant cultivation conditions is therefore a must for future research.

## Supporting information

raw data

## Acknowledgments

The authors wish to thank Frederiek-Maarten Kerckhof for his insights and help with the statistical analysis. The authors also wish to thank Greet Van de Velde and Tim Lacoere for their help with running the IC, HPLC and VFA-GC. Additionally, the authors wish to thank Tim Lacoere for the design of some of the icons used in the graphical abstract.

## CRediT statement

Conceptualization: LD, JVL, JDV, RG; Methodology: LD, JVL, JDV, RG; Formal analysis and investigation: LD, AN, SS, CAF, JD; Writing - original draft preparation: LD; Writing - review and editing: LD, AN, SS, CAF, JVL, JDV, RG; Funding acquisition: JDV, RG; Resources: JDV, RG; Supervision: JVL, JDV, RG

## Availability of data and materials

All data supporting the findings of this study are available within the paper and its Supplementary Information. A copy of the used raw data is provided within the excel file named “20260722_raw-data.xlsx”.

## Ethical approval

Not applicable to this paper

