## Supplementary info for "Cultivation-dependent effects of quorum sensing signals on a lactic acid and chain-elongating bacterium"

Journal: APPLIED MICROBIOLOGY AND BIOTECHNOLOGY

### S1. Experimental conditions and experimental design

#### S1.1 Media composition

The media contained the following per liter: trypticase peptone 5g, yeast extract 0.4g, glucose.$H_2O$ 1.98g, $K_2HPO_4$ 2g, $KH_2PO_4$ 0.23g, $CaCl_2.2H_2O$ 0.25g, NaCl 2.25g, $MgCl_2.6H_2O$ 0.1g, $NH_4Cl$ 2.5g, Wolin modified mineral solution 10mL, resazurin solution (0.1% w/v) 1mL, selenite-tungstate solution 1mL, MES buffer 11.71g and vitamin $K_1$ solution 0.2mL. The modified Wolin mineral solution contained the following per liter: nitrilotriacetic acid 1.5g, $MgSO_4.7H_2O$ 3g, $MnSO_4.H_2O$ 0.5g, NaCl 1g, $FeSO_4.7H_2O$ 0.1g, $CoSO_4.7H_2O$ 0.18g, $CaCl_2.2H_2O$ 0.1g, $ZnSO_4.7H_2O$ 0.18g, $CuSO_4.5H_2O$ 0.01g, $AlK(SO_4)_2.12H_2O$ 0.02g, $H_3BO_3$ 0.01g, $Na_2MoO_4.2H_2O$ 0.01g, $NiCl_2.6H_2O$ 0.03g, $Na_2SeO_3.2H_2O$ 0.3 mg and $Na_2WO_4.2H_2O$ 0.04 mg. The vitamin $K_1$ solution contained 0.1mL vitamin $K_1$ in 20 mL of 95% ethanol. The Wolin vitamin solution contained per liter: biotin 2mg, folic acid 2mg, pyridoxine hydrochloride 10mg, thiamine.HCl 5mg, riboflavin 5mg, nicotinic acid 5mg, calcium D-(+)-pantothenate 5mg, vitamin $B_{12}$ 0.1mg, p-Aminobenzoic acid 5mg and (DL)-alpha-Lipoic acid 5mg. The vitamin solution was prepared as a 10x concentrated stock solution and stored anaerobically, filter-sterilized with a 0.22 µm filter (Macherey-Nagel, Anderlecht, Belgium) in the dark at 4°C. The vitamin solution was before use diluted 10x in sterile, anaerobic $dH_2O$. The selenite-tungstate solution container per liter: NaOH 500mg, Na2SeO3.5$H_2O$ 3mg and Na2WO4.2$H_2O$ 4mg.

In total, twelve conditions were investigated for the pure culture plate experiments, as can be seen in Table S1.

**Table S 1. Conditions tested in this experiment with the full name of the added molecules, the concentration of the quorum sensing molecule in the final media and the corresponding DMSO concentration of the final media.**

| Condition | QS molecule | Manufacturer | Concentration QS molecule | DMSO concentration |
|---|---|---|---|---|
| Control | - | - | - | - |
| DSF | cis-11-Methyl-2-dodecenoic acid | Merck KGaA (Germany) | 1µM | 0.002% |
| C4-HSL | N-Butanoyl-L-homoserine lactone | Merck KGaA (Germany) | 10µM | 0.03% |
| C6-HSL | N-hexanoyl-L-Homoserine lactone | Cayman Chemical (UNITED STATES) | 10µM | 0.04% |
| C8-HSL | N-octanoyl-L-Homoserine lactone | Cayman Chemical (UNITED STATES) | 10µM | 0.04% |
| C10-HSL | N-decanoyl-L-Homoserine lactone | TargetMol (United states) | 10µM | 0.05% |
| C12-HSL | N-dodecanoyl-L-Homoserine lactone | Cayman Chemical (UNITED STATES) | 10µM | 0.06% |
| C14-HSL | N-tetradecanoyl-L-Homoserine lactone | Cayman Chemical (UNITED STATES) | 10µM | 0.06% |
| Indole | Indole | Merck KGaA (Germany) | 500µM | - |
| Farnesol | Farnesol | Merck KGaA (Germany) | 10µM | - |
| EPI | Epinephrine bitartate | Merck KGaA (Germany) | 10µM | - |
| Media | - | - | - | - |

DSF, C4-HSL, C6-HSL, C8-HSL, C10-HSL, C12-HSL and C14-HSL were dissolved in DMSO, filter-sterilized with a 0.22 µm filter (Macherey-Nagel, Anderlecht, Belgium) and stored at -20 °C in brown vials to prevent degradation of the compounds.

### S1.2 Experimental set-up

For the pure culture experiments in plates, each plate contained all 12 conditions (1 control without exogenous quorum sensing molecules, 10 conditions with the exogenous quorum sensing molecules and 1 control without micro-organisms) as seen in Table S 2. For each pure culture experiment, this plate was repeated 3 times for a total of 15 replicates.

**Table S 2. Lay-out of the pure culture experiments in plates. MO stands for micro-organisms, which was either *L. plantarum* or *M. elsdenii* depending on the experiment.**

| | 1 | 2 | 3 | 4 | 5 | 6 | 7 | 8 | 9 | 10 | 11 | 12 |
|---|---|---|---|---|---|---|---|---|---|---|---|---|
| A | | | | | | | | | | | | |
| B | | MO control | MO control | MO control | MO control | MO control | MO + C12-HSL | MO + C12-HSL | MO + C12-HSL | MO + C12-HSL | MO + C12-HSL | |
| C | | MO + DSF | MO + DSF | MO + DSF | MO + DSF | MO + DSF | MO + C14-HSL | MO + C14-HSL | MO + C14-HSL | MO + C14-HSL | MO + C14-HSL | |
| D | | MO + C4-HSL | MO + C4-HSL | MO + C4-HSL | MO + C4-HSL | MO + C4-HSL | MO + indole | MO + indole | MO + indole | MO + indole | MO + indole | |
| E | | MO + C6-HSL | MO + C6-HSL | MO + C6-HSL | MO + C6-HSL | MO + C6-HSL | MO + EPI | MO + EPI | MO + EPI | MO + EPI | MO + EPI | |
| F | | MO + C8-HSL | MO + C8-HSL | MO + C8-HSL | MO + C8-HSL | MO + C8-HSL | MO + farn | MO + farn | MO + farn | MO + farn | MO + farn | |
| G | | MO + C10-HSL | MO + C10-HSL | MO + C10-HSL | MO + C10-HSL | MO + C10-HSL | Blanco media | Blanco media | Blanco media | Blanco media | Blanco media | |
| H | | | | | | | | | | | | |

Additionally, 1 control plate was designed as follows with only DMSO controls according to the concentrations described in Table S 3:

**Table S 3. Lay-out of the controls of the pure culture experiments in plates. MO stands for micro-organisms, which was either *L. plantarum* or *M. elsdenii* depending on the experiment.**

| | 1 | 2 | 3 | 4 | 5 | 6 | 7 | 8 | 9 | 10 | 11 | 12 |
|---|---|---|---|---|---|---|---|---|---|---|---|---|
| A | | | | | | | | | | | | |
| B | | MO | MO + DSF D | MO + C4 D | MO + C6 D | MO + C8 D | MO + C10 D | MO + C12 D | MO + C14 D | media | media +DMSO | |
| C | | MO | MO + DSF D | MO + C4 D | MO + C6 D | MO + C8 D | MO + C10 D | MO + C12 D | MO + C14 D | media | media +DMSO | |
| D | | MO | MO + DSF D | MO + C4 D | MO + C6 D | MO + C8 D | MO + C10 D | MO + C12 D | MO + C14 D | media | media +DMSO | |
| E | | MO | MO + DSF D | MO + C4 D | MO + C6 D | MO + C8 D | MO + C10 D | MO + C12 D | MO + C14 D | media | media +DMSO | |
| F | | MO | MO + DSF D | MO + C4 D | MO + C6 D | MO + C8 D | MO + C10 D | MO + C12 D | MO + C14 D | media | media +DMSO | |
| G | | MO | MO + DSF D | MO + C4 D | MO + C6 D | MO + C8 D | MO + C10 D | MO + C12 D | MO + C14 D | media | media +DMSO | |
| H | | | | | | | | | | | | |

For the co-culture experiments, only conditions were further investigated, namely addition of C8-HSL and addition of C12-HSL. For the co-culture plate experiment, 2 different days were sampled. This was done destructively due to the crystal violet assay. Therefore, 4 plates with 20 replicates per condition each were set-up. After 24 hours, all 4 plates were measured for OD and 2 plates were chosen for further analysis. After 48 hours, the remaining 2 plates were measured for OD and further analyzed. The set-up for this experiment can be found in Table S 4. Since the effects of DMSO were already investigated in the pure culture experiments, no additional plates were set-up to investigate the effect of DMSO on the co-culture.

**Table S 4. Lay-out of the co-culture experiment in plates. LP stands for *L. plantarum*, ME stands for *M. elsdenii*, C8-HSL stands for the addition of exogenous C8-HSL and C12-HSL stands for the addition of exogenous C12-HSL.**

| | 1 | 2 | 3 | 4 | 5 | 6 | 7 | 8 | 9 | 10 | 11 | 12 |
|---|---|---|---|---|---|---|---|---|---|---|---|---|
| A | | | | | | | | | | | | |
| B | | LP + ME | LP + ME | LP + ME | LP + ME | LP + ME | LP + ME | LP + ME | LP + ME | LP + ME | LP + ME | |
| C | | LP+ME +C8-HSL | LP+ME +C8-HSL | LP+ME +C8-HSL | LP+ME +C8-HSL | LP+ME +C8-HSL | LP+ME +C8-HSL | LP+ME +C8-HSL | LP+ME +C8-HSL | LP+ME +C8-HSL | LP+ME +C8-HSL | |
| D | | LP+ME +C12-HSL | LP+ME +C12-HSL | LP+ME +C12-HSL | LP+ME +C12-HSL | LP+ME +C12-HSL | LP+ME +C12-HSL | LP+ME +C12-HSL | LP+ME +C12-HSL | LP+ME +C12-HSL | LP+ME +C12-HSL | |
| E | | LP + ME | LP + ME | LP + ME | LP + ME | LP + ME | LP + ME | LP + ME | LP + ME | LP + ME | LP + ME | |
| F | | LP+ME +C8-HSL | LP+ME +C8-HSL | LP+ME +C8-HSL | LP+ME +C8-HSL | LP+ME +C8-HSL | LP+ME +C8-HSL | LP+ME +C8-HSL | LP+ME +C8-HSL | LP+ME +C8-HSL | LP+ME +C8-HSL | |
| G | | LP+ME +C12-HSL | LP+ME +C12-HSL | LP+ME +C12-HSL | LP+ME +C12-HSL | LP+ME +C12-HSL | LP+ME +C12-HSL | LP+ME +C12-HSL | LP+ME +C12-HSL | LP+ME +C12-HSL | LP+ME +C12-HSL | |
| H | | | | | | | | | | | | |

To ensure inoculation with equal cell numbers of *L. plantarum* and *M. elsdenii*, the cultures were diluted 100x in PBS and measured on the flow cytometer with Sybr Green staining as described in the Material and Methods of the main paper (section 2.4.1). The cell counts for *L. plantarum* and *M. elsdenii* were 25 676 (x100) cells/µL and 7006 (x100) cells/µL, respectively. Therefore, cells are diluted 1:4 *L. plantarum* to *M. elsdenii*.

Prior to the experiments, growth curves were established using the oCelloScope for *L. plantarum*, *M. elsdenii* growing on glucose and *M. elsdenii* growing on lactic acid, as seen in Figure S 1.

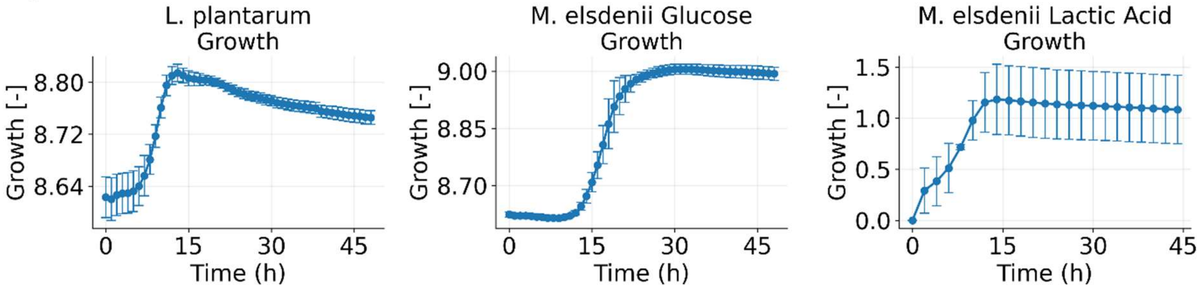

**Figure S 1. Growth curves of *L. plantarum*, *M. elsdenii* growing on glucose and *M. elsdenii* growing on lactic acid. The error bars denote standard deviation (n=3 for *L. plantarum* and *M. elsdenii* growing on glucose, n=5 for *M. elsdenii* growing on lactic acid).**

The growth curves for *L. plantarum* and *M. elsdenii* growing on glucose were obtained during the same experiment, while the growth curve for *M. elsdenii* was obtained during a different experiment. The differences in scale therefore do not reflect differences in growth, but differences in light conditions between the two experiments. For *L. plantarum*, the experiments with exogenous quorum sensing molecules were stopped after 24 hours to ensure limited decay, while for *M. elsdenii*, 48 hours was chosen to account for the slower growth on glucose, while minimizing decay when growing faster on the lactic acid.

For all the experiments, the following number of biological replicates for each of the measured variables were used.

**Table S 5. Description of biological replicates for the growth, biofilm, metabolites, oCelloScope visualization, pH and gas samples taken during the different experiments.**

| Experiment | $n_{growth}$ | $n_{biofilm\ ratio}$ | $n_{Metabolites}$ | $n_{Visualization}$ (full) | $n_{pH}$ | $n_{Gas}$ |
|---|---|---|---|---|---|---|
| Pure culture plate | 15 | 9 | 3 | 15 (5) | - | - |
| Co-culture plate | 40/day | 32/day | 4/day | 40 (20)/day | - | - |
| Serum bottles | 3/12h | - | 3/12h | 3/24h | 3/12h | 3/12h |

#### S1.3 Statistical analysis

Statistical analyses were performed to evaluate whether each treatment differed from the untreated control. Because all treatments were compared with the same untreated control, all treatment-level statistical decisions were interpreted in the context of multiple treatment-versus-control comparisons. The analysis workflow included exploratory plots, formal diagnostic tests, and a decision-based selection of parametric or non-parametric treatment-versus-control tests.

**Diagnostic evaluation of model assumptions**
Before selecting the final treatment-comparison test, each dataset was evaluated visually and statistically. Diagnostic plots included Q-Q plots of model residuals, residuals-versus-fitted plots, and boxplots per condition. These plots were used to assess deviations from normality, heteroscedasticity, outliers, and general distributional differences between conditions.

Residual normality was assessed using the Shapiro-Wilk test on the model residuals. Homogeneity of variance across conditions was assessed using the Brown-Forsythe test, implemented as a median-centered modified Levene test. This was preferred over the classical mean-centered Levene test because it is less sensitive to non-normal distributions. Treatment-specific inference was instead based on the planned treatment-versus-control comparisons described below.

**Treatment-versus-control testing strategy**
The final statistical test was selected based on the diagnostic results and sample size. For all tests, significance was evaluated at $\alpha = 0.05$. When multiple treatments were compared with the same untreated control, adjusted p-values were used.

For OD-based growth measurements and crystal violet biofilm measurements, parametric treatment-versus-control comparisons were used when residual normality was acceptable. Welch's two-sided independent t-tests versus the untreated control with Holm correction were used as the default parametric treatment-versus-control approach for both equal variance and unequal variance because the same reporting strategy was preferred across organisms and datasets.

When both residual normality and variance assumptions were poor, non-parametric treatment-versus-control testing was prioritized. In these cases, and particularly when the Kruskal-Wallis omnibus test indicated an overall condition effect, a Steel-type rank-based permutation test with max-statistic adjustment was used. This test was used instead of independent Mann-Whitney U tests because the experiment involved multiple treatments compared against the same control.

**Steel-type max-statistic permutation test**

The Steel-type test was implemented as a non-parametric treatment-versus-control rank test. For each variable, all observations from the untreated control and all usable treatment groups were pooled and ranked globally. For each treatment, the absolute difference between the treatment mean rank and the control mean rank was calculated. To account for multiple treatment-versus-control comparisons, group labels were randomly permuted while preserving the original group sizes. For each permutation, the maximum absolute treatment-versus-control rank contrast across all treatments was stored, producing a max-statistic null distribution. The adjusted p-value for each treatment was calculated as the proportion of permutations in which the maximum permuted statistic was greater than or equal to the observed statistic for that treatment. This approach controls the family-wise error rate across the set of treatment-versus-control comparisons for a given variable. This Steel-type max-statistic permutation approach was used for datasets where the assumptions of parametric testing were not supported, and for low-replicate datasets such as metabolite measurements. Because metabolite and flow cytometry measurements generally had only three biological replicates per condition, all conclusions from these analyses were interpreted cautiously, regardless of statistical significance. The low sample size limits the power to detect treatment effects and also limits the reliability of formal normality and variance diagnostics.

#### S1.4 Calculations yield

For the element balances, an estimate was made for the biomass based on previously reported yields (Weimer and Moen 2013). Growth of M. elsdenii was reported as 0.079 $\frac{g_{cells}}{g_{glucose}}$ and 0.024 $\frac{g_{cells}}{g_{lactic\ acid}}$. The following equation was used to determine how much carbon went to biomass:

$$\%\ carbon\ to\ biomass\ \left[\frac{Cmol_x}{Cmol_s}\%\right] = \frac{Y_{x/s}\left[\frac{g_x}{g_s}\right]}{MW_x\left[\frac{g_x}{mol_x}\right]} * \frac{MW_s\left[\frac{g_s}{mol_s}\right]}{\#\ of\ C\ in\ s\ \left[\frac{Cmol_s}{mol_s}\right]} * 100\% \qquad \text{Eq. S1}$$

With $x$ the biomass, $s$ the substrate and $MW$ the molecular weight. For the biomass, the generic chemical formula $C_1H_{1.8}O_{0.5}N_{0.2}$ was used, resulting in a molecular weight of 24.6 $\frac{g_x}{mol_x}$. This resulted in 10% of all carbon going to biomass when growing on glucose, and 3% of all carbon going to biomass when growing on lactic acid (including spent supernatant).

### S2. Lactiplantibacillus plantarum pure culture experiments

#### S2.1 *L. plantarum* experiments in plates

The values for the growth and biofilm ratio of *L. plantarum* after 24 hours of growth in a 96 well plate can be found in Table S 6.

**Table S 6. Average measurements for growth ($OD_{620}$) and biofilm ratio ($OD_{540}/OD_{620}$) with the standard deviation and p-value**

| Name condition | Parameter | Replicates (n) | Average ± standard deviation | P-value |
|---|---|---|---|---|
| control | Biofilm ($OD_{540}/OD_{620}$) | 9 | 0.973 ± 0.254 | 0 |
| DSF | Biofilm ($OD_{540}/OD_{620}$) | 9 | 0.83 ± 0.228 | 1 |
| C4-HSL | Biofilm ($OD_{540}/OD_{620}$) | 9 | 0.822 ± 0.165 | 1 |
| C6-HSL | Biofilm ($OD_{540}/OD_{620}$) | 9 | 0.878 ± 0.211 | 1 |
| C8-HSL | Biofilm ($OD_{540}/OD_{620}$) | 9 | 0.941 ± 0.178 | 1 |
| C10-HSL | Biofilm ($OD_{540}/OD_{620}$) | 9 | 0.929 ± 0.19 | 1 |
| C12-HSL | Biofilm ($OD_{540}/OD_{620}$) | 9 | 0.741 ± 0.166 | 0.354001 |
| C14-HSL | Biofilm ($OD_{540}/OD_{620}$) | 9 | 0.952 ± 0.221 | 1 |
| indole | Biofilm ($OD_{540}/OD_{620}$) | 9 | 0.877 ± 0.254 | 1 |
| farnesol | Biofilm ($OD_{540}/OD_{620}$) | 9 | 0.742 ± 0.168 | 0.354001 |
| EPI | Biofilm ($OD_{540}/OD_{620}$) | 9 | 0.73 ± 0.186 | 0.354001 |
| control | Growth ($OD_{620}$) | 15 | 0.226 ± 0.024 | 0 |
| DSF | Growth ($OD_{620}$) | 15 | 0.255 ± 0.026 | 0.0217864 |
| C4-HSL | Growth ($OD_{620}$) | 15 | 0.25 ± 0.024 | 0.0446317 |
| C6-HSL | Growth ($OD_{620}$) | 15 | 0.255 ± 0.027 | 0.027315 |
| C8-HSL | Growth ($OD_{620}$) | 15 | 0.238 ± 0.03 | 0.2618928 |
| C10-HSL | Growth ($OD_{620}$) | 15 | 0.248 ± 0.029 | 0.1017853 |
| C12-HSL | Growth ($OD_{620}$) | 15 | 0.278 ± 0.016 | 3.119E-06 |
| C14-HSL | Growth ($OD_{620}$) | 15 | 0.276 ± 0.01 | 5.388E-06 |
| indole | Growth ($OD_{620}$) | 15 | 0.243 ± 0.019 | 0.1017853 |
| farnesol | Growth ($OD_{620}$) | 15 | 0.271 ± 0.013 | 1.674E-05 |

| | | | | |
|---|---|---|---|---|
| EPI | Growth ($OD_{620}$) | 15 | 0.27 ± 0.011 | 1.999E-05 |

The absolute metabolite production of *L. plantarum* can be found in Figure S 2. Only lactate and formate were detected, with no detected ethanol or acetate as determined by IC and HPLC. Similarly, all glucose was consumed as determined by HPLC.

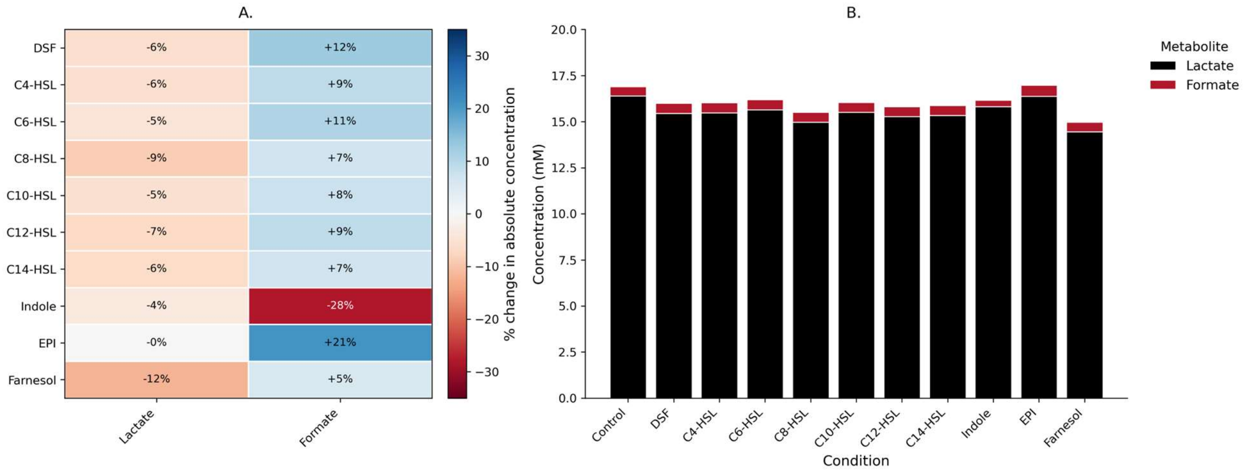

**Figure S 2. Metabolite production of *L. plantarum* after 24 hours with different quorum sensing molecules. * indicates significant changes compared to the untreated control (p < 0.05).**

The values of the detected metabolites, their standard deviation and their adjusted p-value as calculated using the Steel-type max-statistic permutation test ().

**Table S 7. Produced metabolites by *L. plantarum* after 24 hours of growth. The average per condition (n=3) is indicated together with the standard deviation and the adjusted p-value as calculated using the Steel-type max-statistic permutation test.**

| **Name condition** | **Metabolite (mM)** | **Average (n=3) ± standard deviation** | **p value** | **Statistical test** |
|---|---|---|---|---|
| Control | Formate | 0.496 ± 0.0391 | | Steel-type max-statistic permutation test |
| DSF | Formate | 0.558 ± 0.0489 | 0.467653 | Steel-type max-statistic permutation test |
| C4-HSL | Formate | 0.54 ± 0.0507 | 0.705029 | Steel-type max-statistic permutation test |
| C6-HSL | Formate | 0.548 ± 0.0193 | 0.513349 | Steel-type max-statistic permutation test |
| C8-HSL | Formate | 0.53 ± 0.0345 | 0.956204 | Steel-type max-statistic permutation test |
| C10-HSL | Formate | 0.533 ± 0.0324 | 0.89521 | Steel-type max-statistic permutation test |
| C12-HSL | Formate | 0.54 ± 0.0481 | 0.865313 | Steel-type max-statistic permutation test |
| C14-HSL | Formate | 0.531 ± 0.0416 | 0.939406 | Steel-type max-statistic permutation test |

| | | | | |
|---|---|---|---|---|
| Indole | Formate | 0.356 ± 0.31 | 1 | Steel-type max-statistic permutation test |
| EPI | Formate | 0.6 ± 0.0662 | 0.119088 | Steel-type max-statistic permutation test |
| Farnesol | Formate | 0.522 ± 0.0472 | 0.987501 | Steel-type max-statistic permutation test |
| Control | Lactate | 16.4 ± 1.55 | | Steel-type max-statistic permutation test |
| DSF | Lactate | 15.4 ± 0.934 | 0.977402 | Steel-type max-statistic permutation test |
| C4-HSL | Lactate | 15.5 ± 1.41 | 1 | Steel-type max-statistic permutation test |
| C6-HSL | Lactate | 15.7 ± 0.245 | 0.9998 | Steel-type max-statistic permutation test |
| C8-HSL | Lactate | 15 ± 0.981 | 0.640336 | Steel-type max-statistic permutation test |
| C10-HSL | Lactate | 15.5 ± 0.763 | 0.977402 | Steel-type max-statistic permutation test |
| C12-HSL | Lactate | 15.3 ± 0.821 | 0.866513 | Steel-type max-statistic permutation test |
| C14-HSL | Lactate | 15.3 ± 0.852 | 0.945905 | Steel-type max-statistic permutation test |
| Indole | Lactate | 15.8 ± 1.76 | 0.970903 | Steel-type max-statistic permutation test |
| EPI | Lactate | 16.4 ± 1.26 | 1 | Steel-type max-statistic permutation test |
| Farnesol | Lactate | 14.5 ± 2.33 | 0.836516 | Steel-type max-statistic permutation test |

The electron balance for *L. plantarum* can be found in Figure Figure S 3. The electron balance for tested compounds and the control closed between 72% (farnesol) and 82% (control).

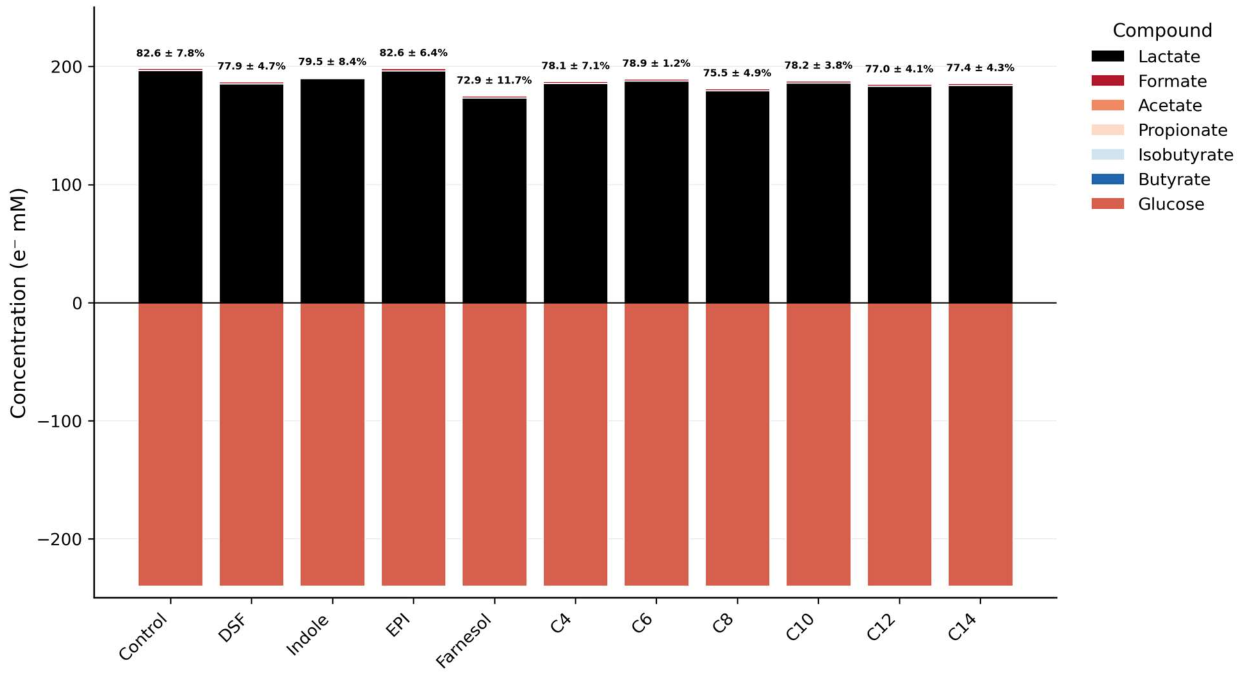

**Figure S 3. Electron balance of *L. plantarum* after 24 hours without (control) or with different quorum sensing molecules. A negative value indicates consumption, while a positive value indicates production.**

To account for potential effects of DMSO, the effect of DMSO on the growth and biofilm formation of *L. plantarum* was also considered (Figure S 4).

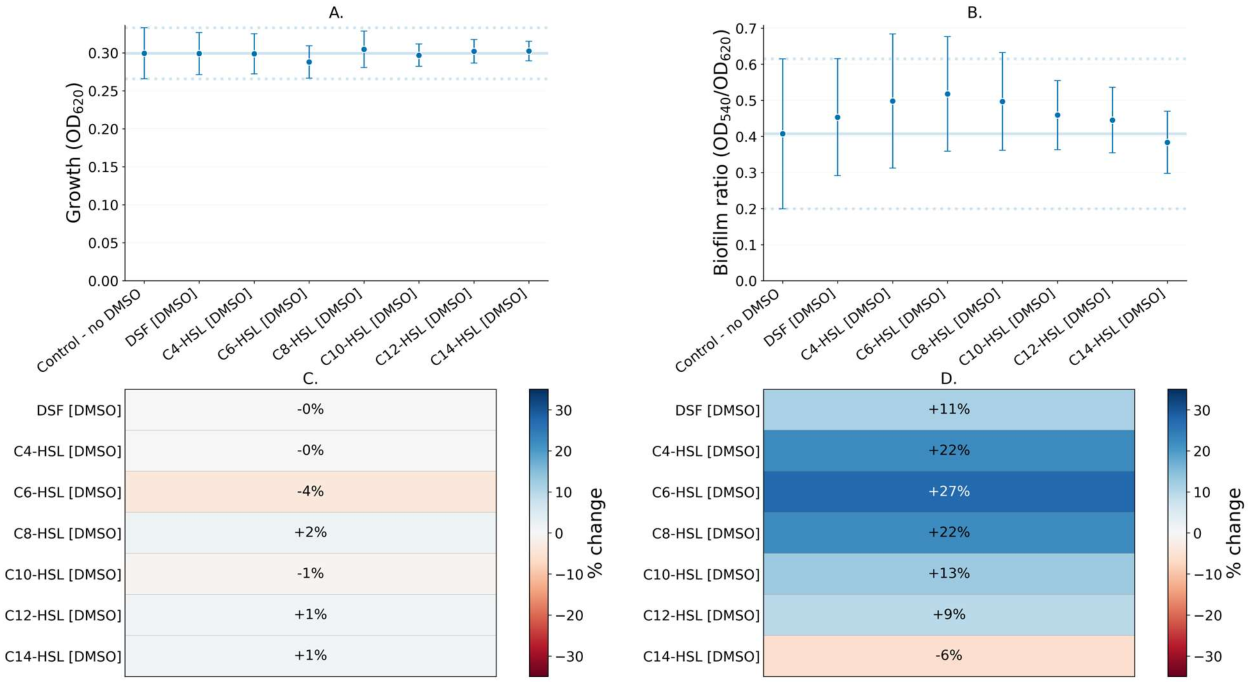

**Figure S 4. Growth (A, n=12) and biofilm ratio (B, n=8) of *L. plantarum* after 24 hours with different DMSO concentrations added corresponding to the quorum sensing conditions or without any DMSO added (control). Error bars indicate standard deviation. The solid line indicates the average value of the condition without exogenous quorum sensing molecules (control), while the dotted lines indicate the mean ± 1 standard deviation. Growth compared to the control (C) and biofilm ratio compared to the control (D) show the change in percent compared to the control. * indicates**

**significant changes compared to the untreated control ($p < 0.05$).**

The values for growth and biofilm for the controls can be found in Table S 8.

**Table S 8. Growth ($OD_{620}$) and biofilm ratio ($OD_{540}/OD_{620}$) of the DMSO controls with the mean values and standard deviation. The p-value is calculated using the test indicated and is considered significant if $p < 0.05$.**

| **variable** | **condition** | **n** | **Mean ± std** | **p-value** | **test** |
|---|---|---|---|---|---|
| Biofilm ($OD_{540}/OD_{620}$) | control | 7 | 0.41 ± 0.21 | - | Steel-type permutation rank test vs control |
| Biofilm ($OD_{540}/OD_{620}$) | DSF | 8 | 0.45 ± 0.16 | 0.9869 | Steel-type permutation rank test vs control |
| Biofilm ($OD_{540}/OD_{620}$) | C4-HSL | 8 | 0.5 ± 0.19 | 0.6725 | Steel-type permutation rank test vs control |
| Biofilm ($OD_{540}/OD_{620}$) | C6-HSL | 8 | 0.52 ± 0.16 | 0.3576 | Steel-type permutation rank test vs control |
| Biofilm ($OD_{540}/OD_{620}$) | C8-HSL | 8 | 0.5 ± 0.14 | 0.5408 | Steel-type permutation rank test vs control |
| Biofilm ($OD_{540}/OD_{620}$) | C10-HSL | 8 | 0.46 ± 0.1 | 0.7481 | Steel-type permutation rank test vs control |
| Biofilm ($OD_{540}/OD_{620}$) | C12-HSL | 8 | 0.45 ± 0.09 | 0.8778 | Steel-type permutation rank test vs control |
| Biofilm ($OD_{540}/OD_{620}$) | C14-HSL | 8 | 0.38 ± 0.09 | 1 | Steel-type permutation rank test vs control |
| Growth ($OD_{620}$) | control | 10 | 0.3 ± 0.03 | - | Steel-type permutation rank test vs control |
| Growth ($OD_{620}$) | DSF | 11 | 0.3 ± 0.03 | 1 | Steel-type permutation rank test vs control |
| Growth ($OD_{620}$) | C4-HSL | 11 | 0.3 ± 0.03 | 1 | Steel-type permutation rank test vs control |
| Growth ($OD_{620}$) | C6-HSL | 11 | 0.29 ± 0.02 | 0.6951 | Steel-type permutation rank test vs control |
| Growth ($OD_{620}$) | C8-HSL | 11 | 0.3 ± 0.02 | 0.9956 | Steel-type permutation rank test vs control |
| Growth ($OD_{620}$) | C10-HSL | 11 | 0.3 ± 0.01 | 0.9976 | Steel-type permutation rank test vs control |
| Growth ($OD_{620}$) | C12-HSL | 11 | 0.3 ± 0.02 | 1 | Steel-type permutation rank test vs control |
| Growth ($OD_{620}$) | C14-HSL | 11 | 0.3 ± 0.01 | 1 | Steel-type permutation rank test vs control |

While growth was largely unaffected, there was some effect on the biofilm ratio, albeit seemingly dose-independent and statistically insignificant. However, the statistical power is considered low due to the small sample size and therefore can only be used as indicative. Nevertheless, due to the lack of apparent effect on the growth and the insignificant change in biofilm ratio, the described effects in this paper were attributed to the quorum sensing molecules and not to DMSO exposure.

#### S2.2 *L. plantarum* serum bottle experiment

For the *L. plantarum* serum bottle experiment, only C8-HSL was investigated due to contamination issues in the C12-HSL condition. The evolution of lactate, formate, growth, pH and $CO_2$ can be seen in Figure S 5.

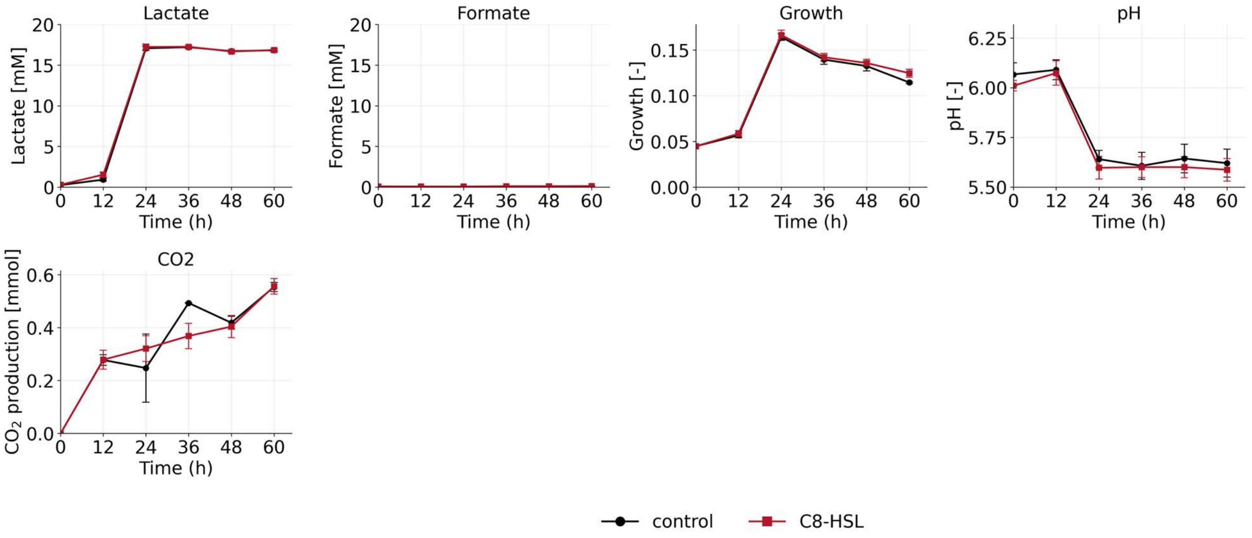

**Figure S 5. Evolution of lactate concentration, formate concentration, growth, pH and $CO_2$ production of *L. plantarum* growing in serum bottles. * indicates significant changes compared to the untreated control ($p < 0.05$).**

Despite the addition of MES buffer, the pH dropped from around 6 to 5.6. Lactate started being produced at 12 hours (0.92 ± 0.15 mM for the control, 1.53 ± 0.27mM for the condition with added C8-HSL) and peaked at 24 hours, with 17.08 ± 0.21mM for the control and 17.23 ± 0.30mM for the condition with added C8-HSL. Growth similarly peaked at 24 hours with an $OD_{620}$ of 0.16 ± 0.00 for the control and 0.17 ±0.00 for the condition with added C8-HSL. The OD then steadily declined until the end of the experiment, where an $OD_{620}$ was measured of 0.11 ± 0.00 for the control and 0.12 ± 0.00 for the condition with added C8-HSL. None of the measured parameters for the condition with C8-HSL was significantly different from the control, which was to be expected as C8-HSL in the pure plate experiments also did not result in a significant difference in either growth or lactate production (Figure S 2, Table S 6 and Table S 7). Glucose was not measured in this experiment.

### S3. Megasphaera elsdenii pure culture experiments

#### S3.1 *M. elsdenii* experiments in plates

The growth and biofilm ratio of *M. elsdenii* after 48 hours in the microtiter plates on the different media with the various quorum sensing molecules can be found in Figure S 6.

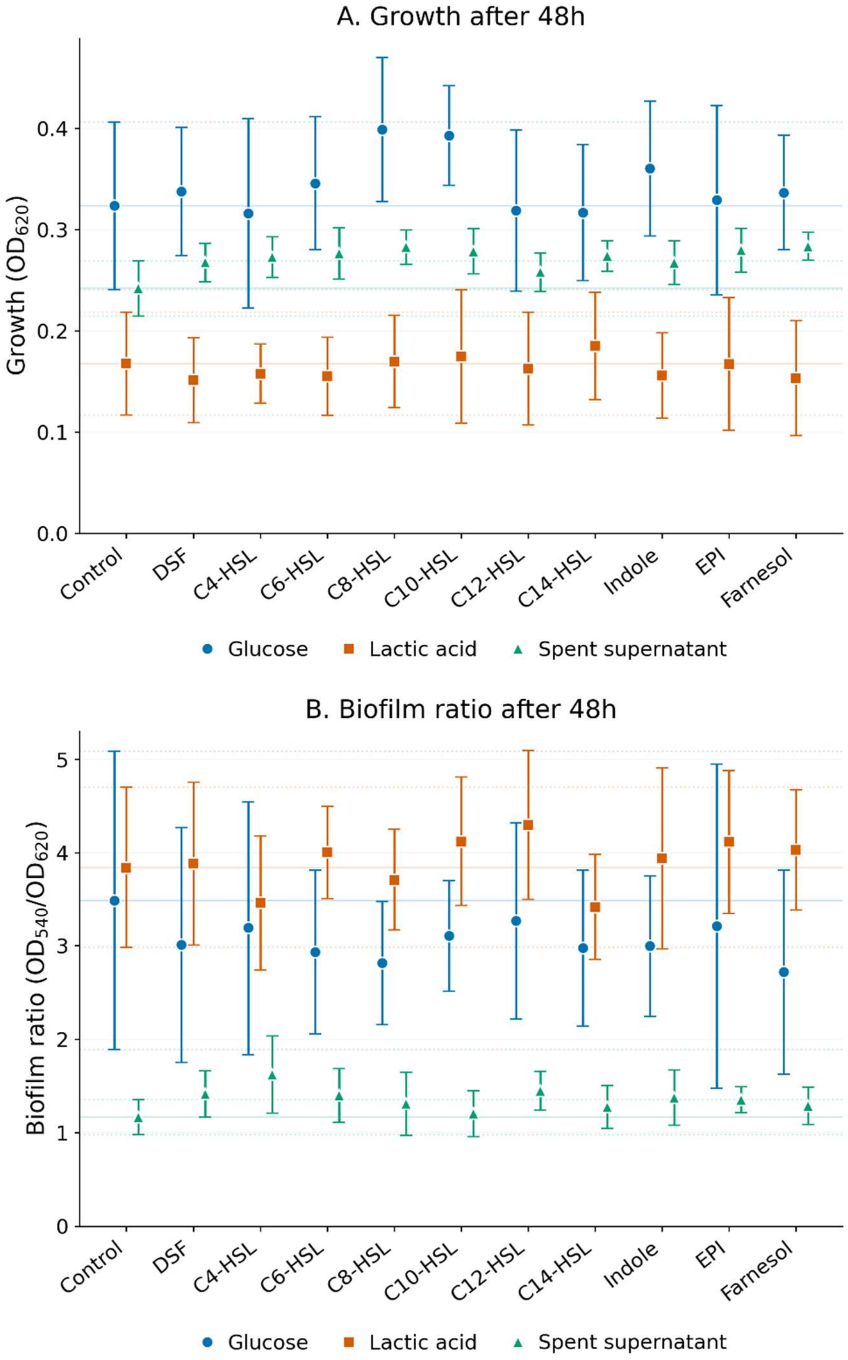

**Figure S 6. Growth (A, n=15) and biofilm ratio (B, n=9) of *M. elsdenii* after 48 hours with different quorum sensing molecules added or without any quorum sensing molecules added (control) growing on glucose (●), lactic acid (■) or spent supernatant (▲). Error bars indicate standard deviation. The full line indicates the average value of the condition without exogenous quorum sensing molecules (control) when growing on glucose (blue), lactic acid (orange) or spent supernatant (green). The dotted lines indicate the confidence interval when growing on glucose (blue), lactic acid**

**(orange) or spent supernatant (green).**

The average values for the growth and biofilm ratio, together with the standard deviation and the chosen statistical test including p-value can be found in Table S 9 (growth) and Table S 10 (biofilm ratio).

| Media | Condition | Mean $OD_{620}$ | Standard deviation | p-value | Selected statistical test |
|---|---|---|---|---|---|
| Glucose | control | 0.3236 | 0.082626 | N/A | Steel-type permutation rank test |
| Glucose | DSF | 0.33782 | 0.063218 | 0.9998 | Steel-type permutation rank test |
| Glucose | C4-HSL | 0.3161 | 0.093555 | 1 | Steel-type permutation rank test |
| Glucose | C6-HSL | 0.345833 | 0.065923 | 0.9884 | Steel-type permutation rank test |
| Glucose | C8-HSL | 0.399033 | 0.071216 | 0.0277 | Steel-type permutation rank test |
| Glucose | C10-HSL | 0.393027 | 0.049241 | 0.0461 | Steel-type permutation rank test |
| Glucose | C12-HSL | 0.31886 | 0.079608 | 1 | Steel-type permutation rank test |
| Glucose | C14-HSL | 0.316853 | 0.067277 | 1 | Steel-type permutation rank test |
| Glucose | Indole | 0.36038 | 0.066544 | 0.7621 | Steel-type permutation rank test |
| Glucose | AI-3 | 0.329133 | 0.093481 | 1 | Steel-type permutation rank test |
| Glucose | Farnesol | 0.336553 | 0.056621 | 1 | Steel-type permutation rank test |
| Lactic acid | control | 0.16768 | 0.050831 | N/A | Steel-type permutation rank test |
| Lactic acid | DSF | 0.151407 | 0.041998 | 0.8461 | Steel-type permutation rank test |
| Lactic acid | C4-HSL | 0.15762 | 0.029108 | 1 | Steel-type permutation rank test |
| Lactic acid | C6-HSL | 0.155073 | 0.038543 | 0.992 | Steel-type permutation rank test |
| Lactic acid | C8-HSL | 0.169687 | 0.045738 | 1 | Steel-type permutation rank test |
| Lactic acid | C10-HSL | 0.174767 | 0.065829 | 1 | Steel-type permutation rank test |
| Lactic acid | C12-HSL | 0.1628 | 0.055636 | 0.9993 | Steel-type permutation rank test |
| Lactic acid | C14-HSL | 0.18508 | 0.052981 | 0.9909 | Steel-type permutation rank test |
| Lactic acid | Indole | 0.15588 | 0.042338 | 0.9699 | Steel-type permutation rank test |
| Lactic acid | AI-3 | 0.1672 | 0.06562 | 0.9905 | Steel-type permutation rank test |
| Lactic acid | Farnesol | 0.153333 | 0.056878 | 0.4299 | Steel-type permutation rank test |
| Spent supernatant | control | 0.241947 | 0.027404 | N/A | Welch t-test with adjusted p-values |
| Spent supernatant | DSF | 0.267547 | 0.019107 | 0.01956 | Welch t-test with adjusted p-values |
| Spent supernatant | C4-HSL | 0.27294 | 0.019999 | 0.006407 | Welch t-test with adjusted p-values |
| Spent supernatant | C6-HSL | 0.276387 | 0.02521 | 0.006407 | Welch t-test with adjusted p-values |
| Spent supernatant | C8-HSL | 0.282733 | 0.017082 | 0.000522 | Welch t-test with adjusted p-values |
| Spent supernatant | C10-HSL | 0.278673 | 0.022122 | 0.002825 | Welch t-test with adjusted p-values |
| Spent supernatant | C12-HSL | 0.257953 | 0.019011 | 0.074904 | Welch t-test with adjusted p-values |
| Spent supernatant | C14-HSL | 0.27396 | 0.01511 | 0.004028 | Welch t-test with adjusted p-values |
| Spent supernatant | Indole | 0.267193 | 0.021555 | 0.01956 | Welch t-test with adjusted p-values |
| Spent supernatant | AI-3 | 0.279693 | 0.021766 | 0.00226 | Welch t-test with adjusted p-values |
| Spent supernatant | Farnesol | 0.283373 | 0.013775 | 0.000367 | Welch t-test with adjusted p-values |

**Table S 9. Mean growth ($OD_{620}$) of *M. elsdenii* growing on various media (with glucose, lactic acid or on the spent supernatant of *L. plantarum*) without exogenous quorum sensing molecules (control) or with exogenous quorum sensing molecules, together with the standard variation, the statistical test and the p-values compared to the control.**

*M. elsdenii* growing on the spent supernatant of *L. plantarum* led to the least amount of biofilm formed (normalized for growth), followed by *M. elsdenii* growing on glucose (Figure S 6). *M. elsdenii* growing on lactic acid produced the most biofilm (normalized for growth) of all conditions. This is also visible in their respective growth morphologies, which are significantly different when growing on glucose (Figure S 7A) versus growing on lactic acid (Figure S 7B) and on the spent supernatant (Figure S 7C).

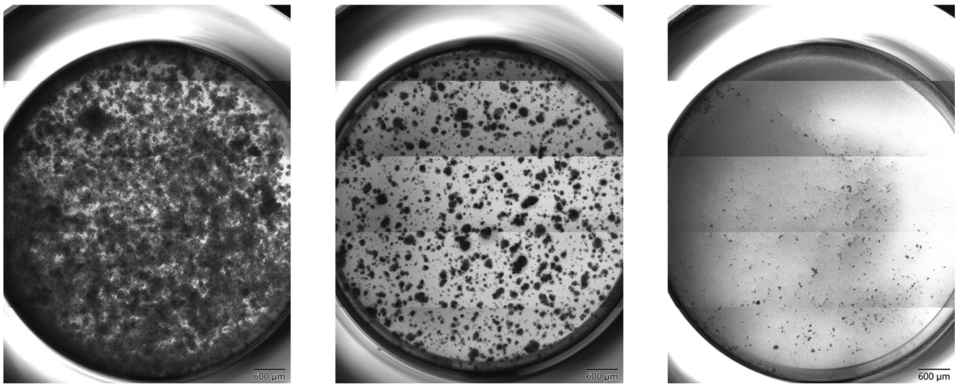

**Figure S 7. Morphology of the *M. elsdenii* control (without exogenous quorum sensing molecules) after 48 hours growing on glucose (A), on lactic acid (B) and growing on *L. plantarum* supernatant (C).**

While growing on glucose, *M. elsdenii* grew in a more concentrated manner, forming dense colonies that adhered strongly to the plate as observed during the experiment. When growing on lactic acid, the culture showed a pronounced clustering with markedly less coverage of the well compared to the glucose condition. This can be partially attributed to the 2-fold decrease in growth (Figure 3), although the clustering itself can be a cause for differences in OD measurements themselves. While some of these colonies were still visible when growing on the *L. plantarum* supernatant, the size and number of clusters decreased both in number and in adherence behavior, with easier detachment from the plate and seemingly more planktonic growth compared to attached biofilm as observed during the experiment.

The values for the biofilm ratio of *M. elsdenii* growing on the different media without exogenous quorum sensing molecules or with added quorum sensing molecules can be found in Table S 10.

| **Media** | **Condition** | **Mean biofilm ratio ($OD_{540}$ /$OD_{620}$)** | **Standard deviation** | **p-value** | **Selected statistical test** |
|---|---|---|---|---|---|
| Glucose | control | 3.488722 | 1.600468 | N/A | Steel-type permutation rank test |
| Glucose | DSF | 3.010133 | 1.257379 | 0.9985 | Steel-type permutation rank test |
| Glucose | C4-HSL | 3.190938 | 1.356485 | 1 | Steel-type permutation rank test |
| Glucose | C6-HSL | 2.933786 | 0.878393 | 0.9999 | Steel-type permutation rank test |
| Glucose | C8-HSL | 2.816614 | 0.663633 | 0.996 | Steel-type permutation rank test |
| Glucose | C10-HSL | 3.105779 | 0.59267 | 1 | Steel-type permutation rank test |
| Glucose | C12-HSL | 3.267652 | 1.053823 | 1 | Steel-type permutation rank test |
| Glucose | C14-HSL | 2.975391 | 0.835867 | 1 | Steel-type permutation rank test |
| Glucose | Indole | 2.996506 | 0.75441 | 1 | Steel-type permutation rank test |
| Glucose | EPI | 3.209995 | 1.738055 | 0.9977 | Steel-type permutation rank test |
| Glucose | Farnesol | 2.718018 | 1.093989 | 0.7502 | Steel-type permutation rank test |
| Lactic acid | control | 3.840474 | 0.861172 | N/A | Welch t-test with adjusted p-values |
| Lactic acid | DSF | 3.880888 | 0.873212 | 1 | Welch t-test with adjusted p-values |

| | | | | | |
|---|---|---|---|---|---|
| Lactic acid | C4-HSL | 3.461817 | 0.72059 | 1 | Welch t-test with adjusted p-values |
| Lactic acid | C6-HSL | 4.002732 | 0.494431 | 1 | Welch t-test with adjusted p-values |
| Lactic acid | C8-HSL | 3.707678 | 0.542654 | 1 | Welch t-test with adjusted p-values |
| Lactic acid | C10-HSL | 4.122961 | 0.688508 | 1 | Welch t-test with adjusted p-values |
| Lactic acid | C12-HSL | 4.297536 | 0.797935 | 1 | Welch t-test with adjusted p-values |
| Lactic acid | C14-HSL | 3.417874 | 0.562438 | 1 | Welch t-test with adjusted p-values |
| Lactic acid | Indole | 3.938914 | 0.970354 | 1 | Welch t-test with adjusted p-values |
| Lactic acid | EPI | 4.114496 | 0.765534 | 1 | Welch t-test with adjusted p-values |
| Lactic acid | Farnesol | 4.031011 | 0.644796 | 1 | Welch t-test with adjusted p-values |
| Spent supernatant | control | 1.165112 | 0.186956 | N/A | Welch t-test with adjusted p-values |
| Spent supernatant | DSF | 1.413688 | 0.248089 | 0.266922 | Welch t-test with adjusted p-values |
| Spent supernatant | C4-HSL | 1.623221 | 0.412224 | 0.103716 | Welch t-test with adjusted p-values |
| Spent supernatant | C6-HSL | 1.398911 | 0.2885 | 0.390778 | Welch t-test with adjusted p-values |
| Spent supernatant | C8-HSL | 1.307786 | 0.339379 | 0.889224 | Welch t-test with adjusted p-values |
| Spent supernatant | C10-HSL | 1.203882 | 0.245745 | 0.889224 | Welch t-test with adjusted p-values |
| Spent supernatant | C12-HSL | 1.447823 | 0.209748 | 0.101728 | Welch t-test with adjusted p-values |
| Spent supernatant | C14-HSL | 1.273268 | 0.229434 | 0.889224 | Welch t-test with adjusted p-values |
| Spent supernatant | Indole | 1.374445 | 0.294195 | 0.494834 | Welch t-test with adjusted p-values |
| Spent supernatant | EPI | 1.353086 | 0.13764 | 0.266922 | Welch t-test with adjusted p-values |
| Spent supernatant | Farnesol | 1.287928 | 0.199728 | 0.841059 | Welch t-test with adjusted p-values |

**Table S 10. Mean biofilm ratio of *M. elsdenii* growing on various media (with glucose, lactic acid or on the spent supernatant of *L. plantarum*) without exogenous quorum sensing molecules (control) or with exogenous quorum sensing molecules, together with the standard variation, the statistical test and the p-values compared to the control.**

The metabolite composition can be found in Figure S 8.

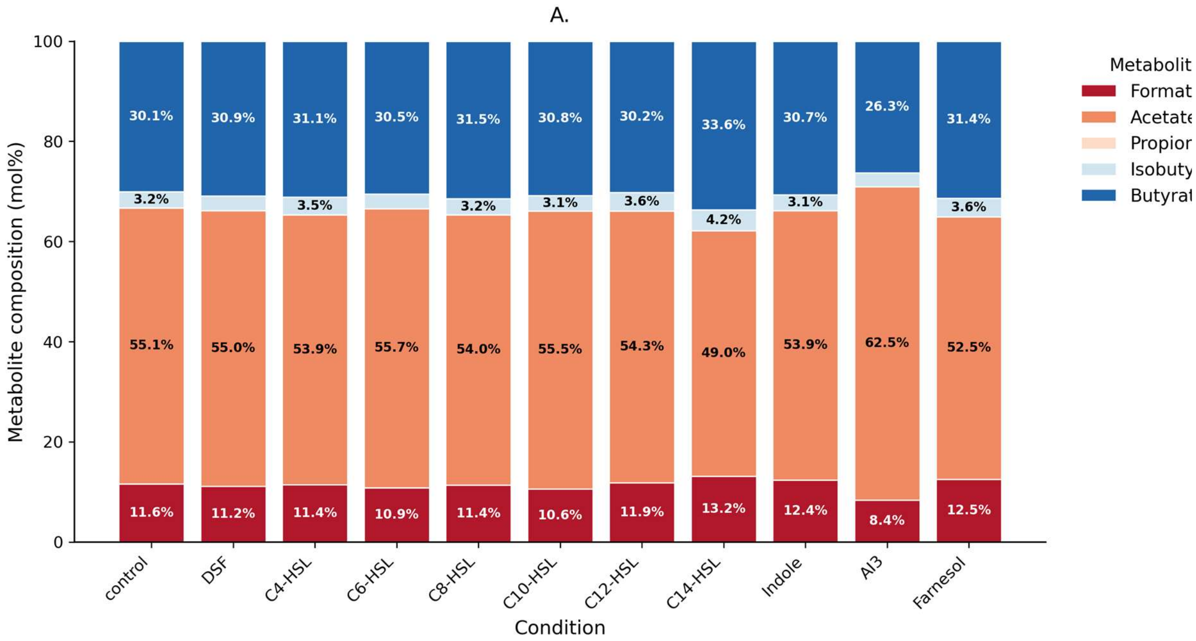

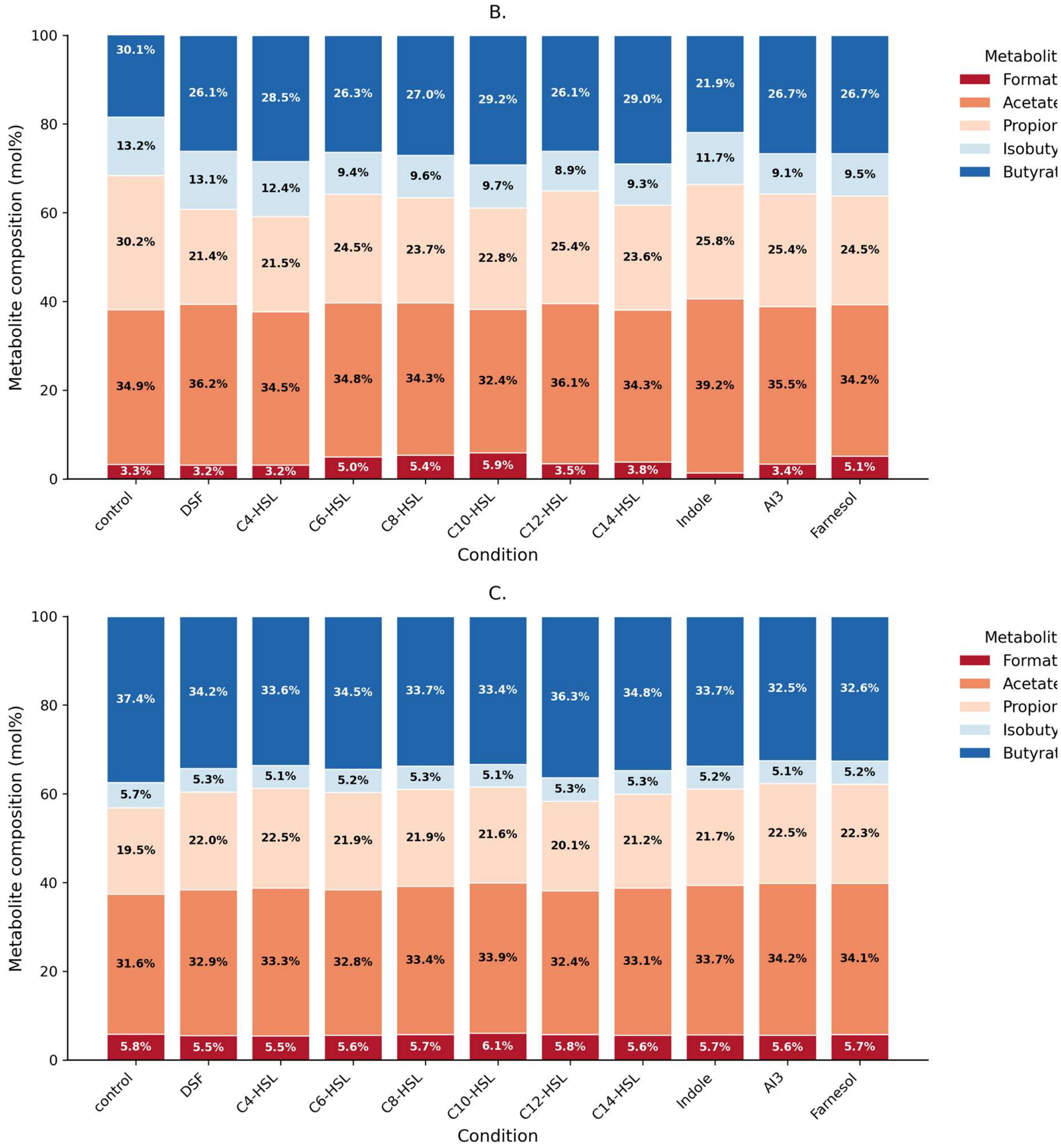

**Figure S 8. Metabolite composition profiles in mol% after 48 hours of *M. elsdenii* growing on glucose (A), lactic acid (B) and spent supernatant of *L. plantarum* (C). Values in the figure indicate the average mol% (n=3) of that compound.**

The metabolite data for the plate experiment can be found in Table S 11 (glucose), Table S 12 (lactic acid) and Table S 13 (spent supernatant of *L. plantarum*).

**Table S 11. Metabolites of *M. elsdenii* growing on glucose for 48 hours. Values are represented as means ± standard deviation (p-value as determined using Steel-type max permutation test).**

| Condition | Formate (mM) | Acetate (mM) | Propionate (mM) | Isobutyrate (mM) | Butyrate (mM) |
|---|---|---|---|---|---|
| control | 1.06 ± 0.12 | 5.04 ± 0.66 | 0 ± 0 | 0.29 ± 0.02 | 2.75 ± 0.29 |
| DSF | 1.05 ± 0.07 (p = 1) | 5.23 ± 0.5 (p = 1) | 0 ± 0 (p = 1) | 0.28 ± 0.06 (p = 1) | 2.95 ± 0.56 (p = 1) |
| C4-HSL | 1.1 ± 0.04 (p = 0.97) | 5.2 ± 0.57 (p = 1) | 0 ± 0 (p = 1) | 0.34 ± 0.05 (p = 0.81) | 3.02 ± 0.48 (p = 1) |

| | | | | | |
|---|---|---|---|---|---|
| C6-HSL | 0.99 ± 0.01 (p = 0.99) | 5.14 ± 0.6 (p = 1) | 0 ± 0 (p = 1) | 0.27 ± 0.04 (p = 1) | 2.83 ± 0.55 (p = 1) |
| C8-HSL | 1.07 ± 0.09 (p = 1) | 5.08 ± 0.16 (p = 1) | 0 ± 0 (p = 1) | 0.3 ± 0.03 (p = 1) | 2.97 ± 0.39 (p = 1) |
| C10-HSL | 0.99 ± 0.06 (p = 0.86) | 5.19 ± 0.44 (p = 1) | 0 ± 0 (p = 1) | 0.29 ± 0 (p = 1) | 2.88 ± 0.3 (p = 1) |
| C12-HSL | 1.01 ± 0.04 (p = 1) | 4.8 ± 1.13 (p = 1) | 0 ± 0 (p = 1) | 0.31 ± 0.05 (p = 1) | 2.69 ± 0.82 (p = 1) |
| C14-HSL | 0.96 ± 0.06 (p = 0.75) | 4.03 ± 1.95 (p = 0.99) | 0 ± 0 (p = 1) | 0.31 ± 0.08 (p = 1) | 2.55 ± 0.54 (p = 1) |
| Indole | 1.03 ± 0.08 (p = 1) | 4.56 ± 0.66 (p = 0.95) | 0 ± 0 (p = 1) | 0.26 ± 0.04 (p = 0.99) | 2.61 ± 0.5 (p = 1) |
| AI-3 | 0.66 ± 0.57 (p = 0.69) | 6.43 ± 3.44 (p = 1) | 0 ± 0 (p = 1) | 0.25 ± 0.04 (p = 0.92) | 2.49 ± 0.45 (p = 1) |
| Farnesol | 1.01 ± 0.03 (p = 1) | 4.34 ± 0.87 (p = 0.79) | 0 ± 0 (p = 1) | 0.29 ± 0.07 (p = 1) | 2.62 ± 0.74 (p = 1) |

**Table S 12. Metabolites of *M. elsdenii* growing on lactic acid for 48 hours. Values are represented as means ± standard deviation (p-value as determined using Steel-type max permutation test).**

| Condition | Formate (mM) | Acetate (mM) | Propionate (mM) | Isobutyrate (mM) | Butyrate (mM) |
|---|---|---|---|---|---|
| control | 0.21 ± 0.18 (p = 0) | 2.08 ± 0.53 (p = 0) | 1.71 ± 0.42 (p = 0) | 0.74 ± 0.04 (p = 0) | 1.79 ± 0.8 (p = 0) |
| DSF | 0.25 ± 0.22 (p = 1) | 2.19 ± 0.72 (p = 1) | 1.39 ± 0.71 (p = 1) | 0.76 ± 0.21 (p = 1) | 1.67 ± 0.79 (p = 0.98) |
| C4-HSL | 0.25 ± 0.22 (p = 1) | 2.1 ± 0.83 (p = 1) | 1.41 ± 0.8 (p = 1) | 0.67 ± 0.1 (p = 0.96) | 1.78 ± 0.81 (p = 1) |
| C6-HSL | 0.38 ± 0.07 (p = 0.81) | 2.66 ± 0.29 (p = 0.66) | 1.88 ± 0.33 (p = 1) | 0.72 ± 0.05 (p = 1) | 2 ± 0.04 (p = 0.98) |
| C8-HSL | 0.42 ± 0.09 (p = 0.35) | 2.68 ± 0.12 (p = 0.69) | 1.86 ± 0.2 (p = 1) | 0.75 ± 0.02 (p = 1) | 2.11 ± 0.11 (p = 1) |
| C10-HSL | 0.44 ± 0.03 (p = 0.16) | 2.41 ± 0.16 (p = 1) | 1.7 ± 0.18 (p = 1) | 0.72 ± 0.01 (p = 0.97) | 2.17 ± 0.09 (p = 1) |
| C12-HSL | 0.28 ± 0.25 (p = 0.92) | 2.47 ± 0.75 (p = 0.85) | 1.79 ± 0.74 (p = 1) | 0.62 ± 0.23 (p = 1) | 1.74 ± 0.36 (p = 0.72) |
| C14-HSL | 0.29 ± 0.25 (p = 0.92) | 2.18 ± 0.53 (p = 1) | 1.52 ± 0.44 (p = 1) | 0.61 ± 0.22 (p = 0.87) | 1.85 ± 0.47 (p = 1) |
| Indole | 0.1 ± 0.17 (p = 1) | 1.86 ± 0.33 (p = 1) | 1.24 ± 0.26 (p = 0.6) | 0.57 ± 0.16 (p = 0.32) | 1.18 ± 0.85 (p = 0.64) |
| AI-3 | 0.27 ± 0.24 (p = 0.96) | 2.45 ± 0.54 (p = 0.87) | 1.79 ± 0.59 (p = 1) | 0.64 ± 0.21 (p = 1) | 1.83 ± 0.37 (p = 0.99) |
| Farnesol | 0.38 ± 0.06 (p = 0.81) | 2.54 ± 0.17 (p = 0.95) | 1.83 ± 0.21 (p = 1) | 0.7 ± 0.05 (p = 0.99) | 1.98 ± 0.14 (p = 1) |

**Table S 13. Metabolites of *M. elsdenii* growing on the spent supernatant of *L. plantarum* for 48 hours. Values are represented as means ± standard deviation (p-value as determined using Steel-type max permutation test).**

| Condition | Formate (mM) | Acetate (mM) | Propionate (mM) | Isobutyrate (mM) | Butyrate (mM) |
|---|---|---|---|---|---|
| control | 0.59 ± 0.02 | 3.2 ± 0.16 | 1.98 ± 0.05 | 0.58 ± 0.03 | 3.79 ± 0.13 |
| DSF | 0.58 ± 0.02 (p = 1) | 3.42 ± 0.27 (p = 0.77) | 2.29 ± 0.23 (p = 0.41) | 0.56 ± 0.03 (p = 1) | 3.56 ± 0.28 (p = 0.89) |

| | | | | | |
|---|---|---|---|---|---|
| C4-HSL | 0.56 ± 0.04 (p = 0.99) | 3.44 ± 0.32 (p = 0.94) | 2.33 ± 0.3 (p = 0.41) | 0.53 ± 0.05 (p = 0.93) | 3.47 ± 0.27 (p = 0.68) |
| C6-HSL | 0.58 ± 0.05 (p = 1) | 3.43 ± 0.1 (p = 0.77) | 2.29 ± 0.07 (p = 0.26) | 0.55 ± 0.02 (p = 0.98) | 3.61 ± 0.23 (p = 0.98) |
| C8-HSL | 0.61 ± 0.03 (p = 1) | 3.57 ± 0.32 (p = 0.57) | 2.33 ± 0.24 (p = 0.25) | 0.57 ± 0.07 (p = 0.95) | 3.59 ± 0.24 (p = 0.98) |
| C10-HSL | 0.57 ± 0.07 (p = 1) | 3.18 ± 0.35 (p = 1) | 2.04 ± 0.29 (p = 1) | 0.48 ± 0.06 (p = 0.1) | 3.13 ± 0.29 (p = 0.04) |
| C12-HSL | 0.51 ± 0.01 (p = 0.15) | 2.88 ± 0.19 (p = 0.94) | 1.79 ± 0.11 (p = 1) | 0.48 ± 0.07 (p = 0.12) | 3.24 ± 0.42 (p = 0.2) |
| C14-HSL | 0.55 ± 0.05 (p = 0.74) | 3.22 ± 0.2 (p = 1) | 2.06 ± 0.19 (p = 1) | 0.52 ± 0.09 (p = 0.81) | 3.4 ± 0.5 (p = 0.66) |
| Indole | 0.57 ± 0.04 (p = 1) | 3.35 ± 0.09 (p = 1) | 2.15 ± 0.11 (p = 0.9) | 0.52 ± 0.04 (p = 0.37) | 3.35 ± 0.3 (p = 0.34) |
| AI-3 | 0.57 ± 0.03 (p = 1) | 3.47 ± 0.24 (p = 0.77) | 2.29 ± 0.29 (p = 0.48) | 0.52 ± 0.08 (p = 0.93) | 3.31 ± 0.42 (p = 0.37) |
| Farnesol | 0.58 ± 0.04 (p = 1) | 3.45 ± 0.1 (p = 0.67) | 2.26 ± 0.11 (p = 0.48) | 0.53 ± 0.02 (p = 0.56) | 3.29 ± 0.21 (p = 0.19) |

The electron balances for the metabolites can be found in Figure S 9A (growth on glucose), Figure S 9B (growth on lactic acid) and Figure S 9C (growth on spent *L. plantarum* supernatant).

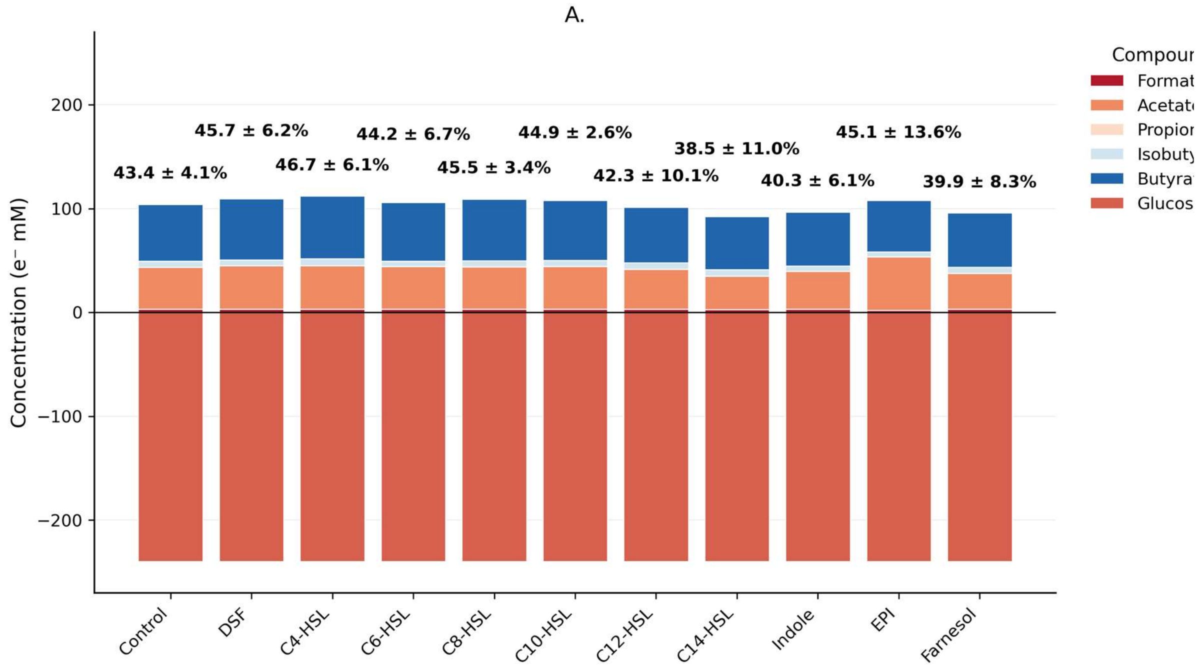

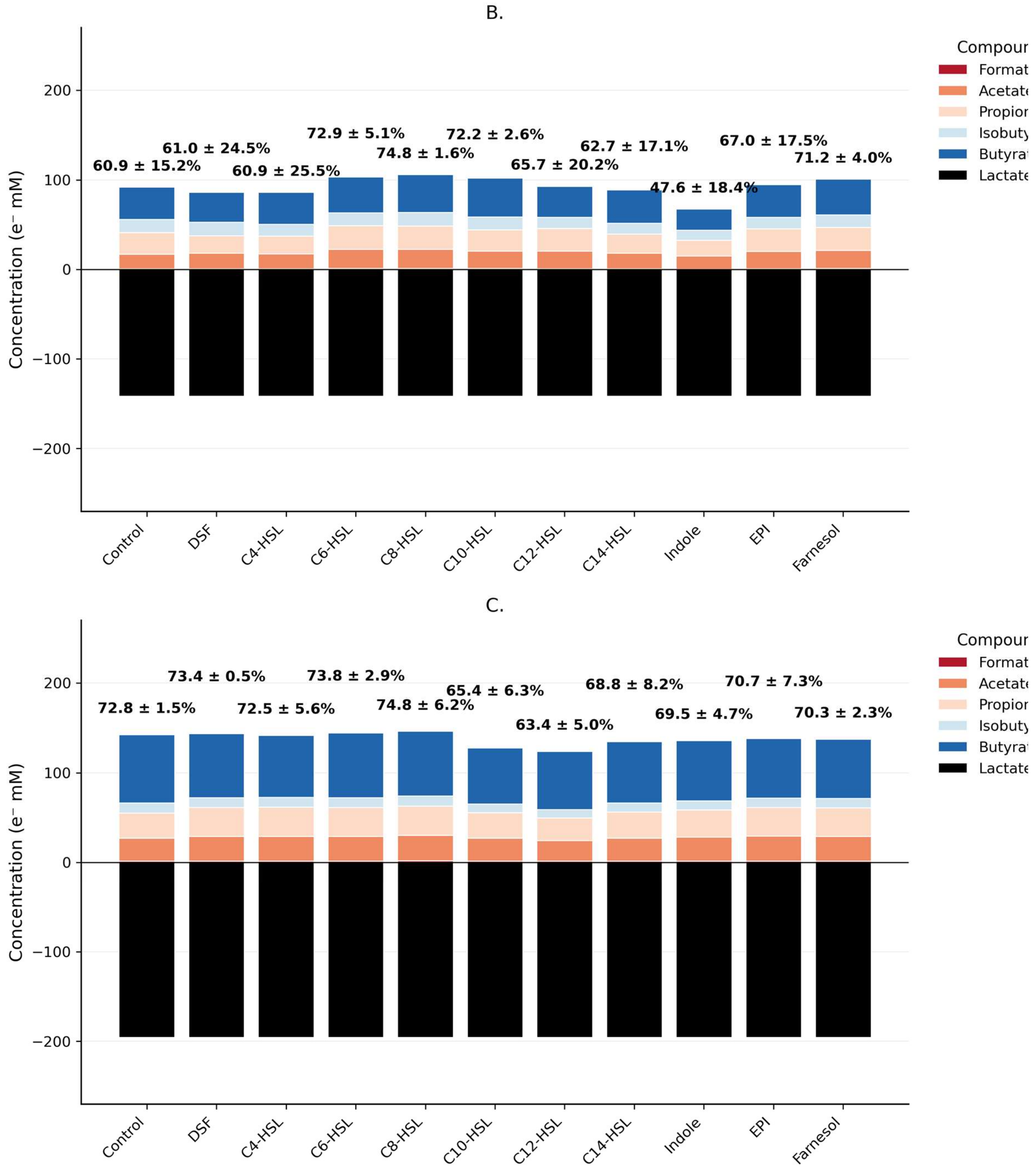

**Figure S 9. Electron balances of *M. elsdenii* grown for 48 hours on glucose (A), lactic acid (B) and spent supernatant of *L. plantarum* (C) without exogenous quorum sensing molecules (control) and with added quorum sensing molecules. Positive values indicate production, while negative values indicate consumption. Values above indicate the degree of closure (%) ± the standard deviation.**

As can be seen in the figures, the electron balance for the spent supernatant closes the best with a range of 63% (C12-HSL) to 75% (C8-HSL). Combined with the produced $H_2$ and the produced biomass, this indicates that most of the metabolites are accounted for. This is not the case for either growth on glucose and growth on lactic acid, which both have a range of 40% to 48%, indicating a likely other metabolic product produced that wasn't measured. For growth on glucose, this product is likely to be hexanoate, while on lactic acid this product is likely to be valerate. Therefore, a calculation as described in section 2.7 was made to estimate the remaining unmeasured compounds, as seen in Figure S 10.

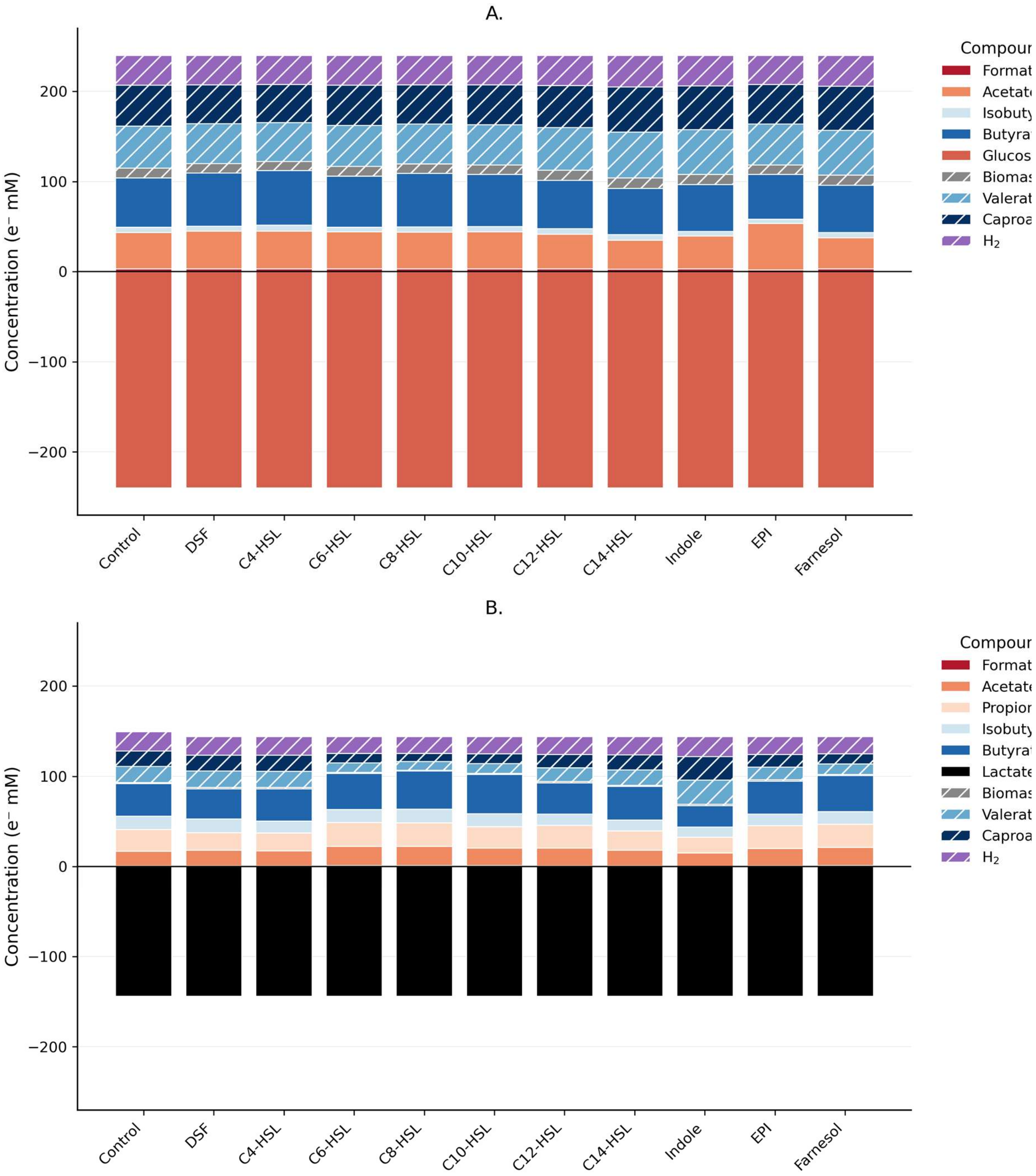

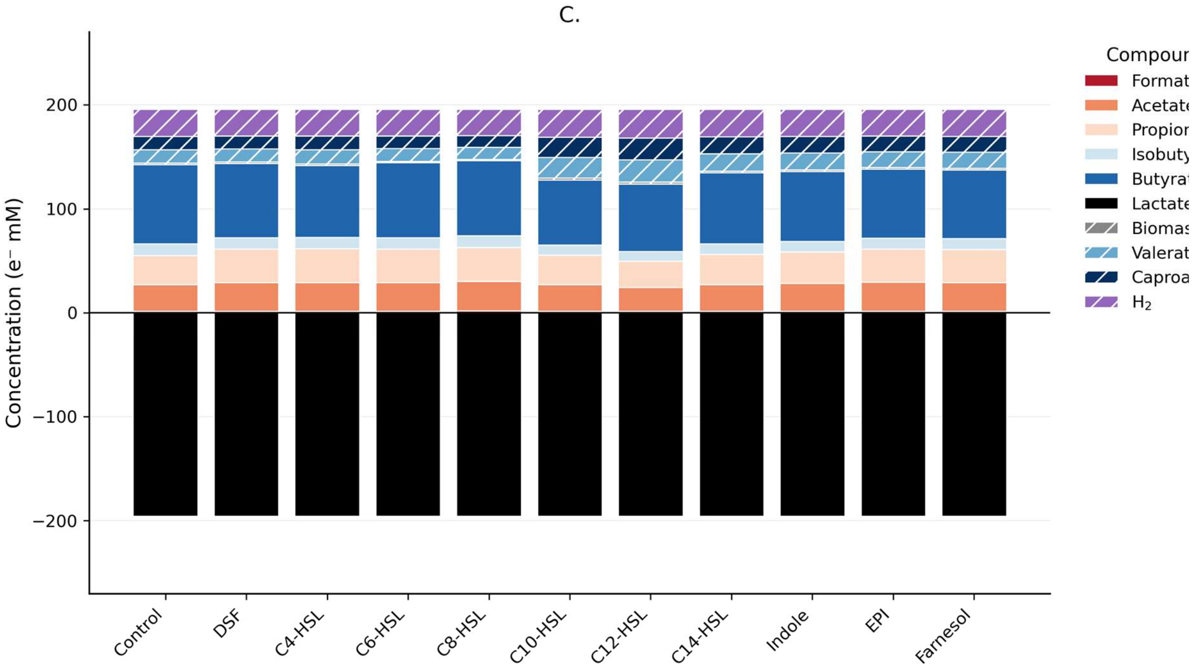

**Figure S 10. Electron balances of *M. elsdenii* grown for 48 hours on glucose (A), lactic acid (B) and spent supernatant of *L. plantarum* (C) without exogenous quorum sensing molecules (control) and with added quorum sensing molecules. Positive values indicate production, while negative values indicate consumption. Calculated values are indicated with a cross-hatch pattern.**

Table S 14, Table S 15 and Table S 16 show the calculated values for biomass, valerate, caproate, $CO_2$ and $H_2$ for M. elsdenii growing on glucose (Table S 14), lactic acid (Table S 15) and spent supernatant (Table S 16).

**Table S 14. Calculated values for biomass, valerate, caproate, $CO_2$ and $H_2$ for *M. elsdenii* growing on glucose for 48 hours in a microtiter plate**

| Condition | Biomass (mM) | Valerate (mM) | Caproate (mM) | $CO_2$ (mM) | $H_2$ (mM) |
|---|---|---|---|---|---|
| control | 2.570825 | 1.79003 | 1.424186 | 16.94043 | 16.42409 |
| DSF | 2.488966 | 1.697345 | 1.350461 | 16.74972 | 16.23542 |
| C4-HSL | 2.453986 | 1.655825 | 1.317413 | 16.68706 | 16.15046 |
| C6-HSL | 2.54229 | 1.754247 | 1.395755 | 16.90772 | 16.42264 |
| C8-HSL | 2.497957 | 1.701139 | 1.353476 | 16.8331 | 16.31136 |
| C10-HSL | 2.51719 | 1.724807 | 1.372339 | 16.8592 | 16.37624 |
| C12-HSL | 2.616179 | 1.828211 | 1.454589 | 17.17462 | 16.68007 |
| C14-HSL | 2.766758 | 1.965947 | 1.564201 | 17.84525 | 17.37612 |
| Indole | 2.688769 | 1.905952 | 1.516426 | 17.38722 | 16.88616 |
| EPI | 2.487244 | 1.734097 | 1.379859 | 16.36651 | 16.04653 |
| Farnesol | 2.708502 | 1.915747 | 1.524235 | 17.55572 | 17.06311 |

**Table S 15. Calculated values for biomass, valerate, caproate, $CO_2$ and $H_2$ for *M. elsdenii* growing on lactic acid for 48 hours in a microtiter plate**

| Condition | Biomass (mM) | Valerate (mM) | Caproate (mM) | CO2 (mM) | H2 (mM) |
|---|---|---|---|---|---|

| | | | | | |
|---|---|---|---|---|---|
| control | 0.343201 | 0.638178 | 0.506885 | 10.62208 | 10.51938 |
| DSF | 0.337331 | 0.65494 | 0.520207 | 10.17006 | 10.04857 |
| C4-HSL | 0.338772 | 0.656519 | 0.521461 | 10.22543 | 10.10331 |
| C6-HSL | 0.263388 | 0.370551 | 0.29418 | 9.316917 | 9.127149 |
| C8-HSL | 0.2523 | 0.323304 | 0.256633 | 9.233886 | 9.024927 |
| C10-HSL | 0.269447 | 0.38271 | 0.303821 | 9.495827 | 9.276764 |
| C12-HSL | 0.306881 | 0.547834 | 0.435078 | 9.720995 | 9.579925 |
| C14-HSL | 0.327142 | 0.615196 | 0.488607 | 10.05802 | 9.91556 |
| Indole | 0.417771 | 0.987539 | 0.784644 | 10.87082 | 10.8244 |
| EPI | 0.299385 | 0.510355 | 0.405297 | 9.718949 | 9.584583 |
| Farnesol | 0.274043 | 0.410767 | 0.326131 | 9.44738 | 9.257253 |

**Table S 16. Calculated values for biomass, valerate, caproate, $CO_2$ and $H_2$ for *M. elsdenii* growing on spent supernatant of *L. plantarum* for 48 hours in a microtiter plate**

| Condition | Biomass (mM) | Valerate (mM) | Caproate (mM) | CO2 (mM) | H2 (mM) |
|---|---|---|---|---|---|
| control | 0.369792 | 0.495893 | 0.393656 | 13.31875 | 13.02621 |
| DSF | 0.36273 | 0.485815 | 0.385656 | 13.07035 | 12.78414 |
| C4-HSL | 0.369274 | 0.517223 | 0.410613 | 13.0849 | 12.8055 |
| C6-HSL | 0.359563 | 0.472313 | 0.374927 | 13.04667 | 12.75623 |
| C8-HSL | 0.350415 | 0.442153 | 0.350962 | 12.89204 | 12.58904 |
| C10-HSL | 0.428223 | 0.765388 | 0.607802 | 13.55587 | 13.2724 |
| C12-HSL | 0.447944 | 0.824291 | 0.654625 | 13.94892 | 13.69546 |
| C14-HSL | 0.401202 | 0.641199 | 0.50912 | 13.44196 | 13.17022 |
| Indole | 0.394202 | 0.621155 | 0.493189 | 13.29398 | 13.01322 |
| EPI | 0.38359 | 0.585084 | 0.464528 | 13.12518 | 12.84077 |
| Farnesol | 0.386975 | 0.599024 | 0.475601 | 13.15526 | 12.86616 |

To account for potential effects of DMSO on the growth and biofilm formation of *M. elsdenii* in the different media, controls with the same DMSO concentration as the conditions (Table S 1) were set-up as seen in Figure S 11.

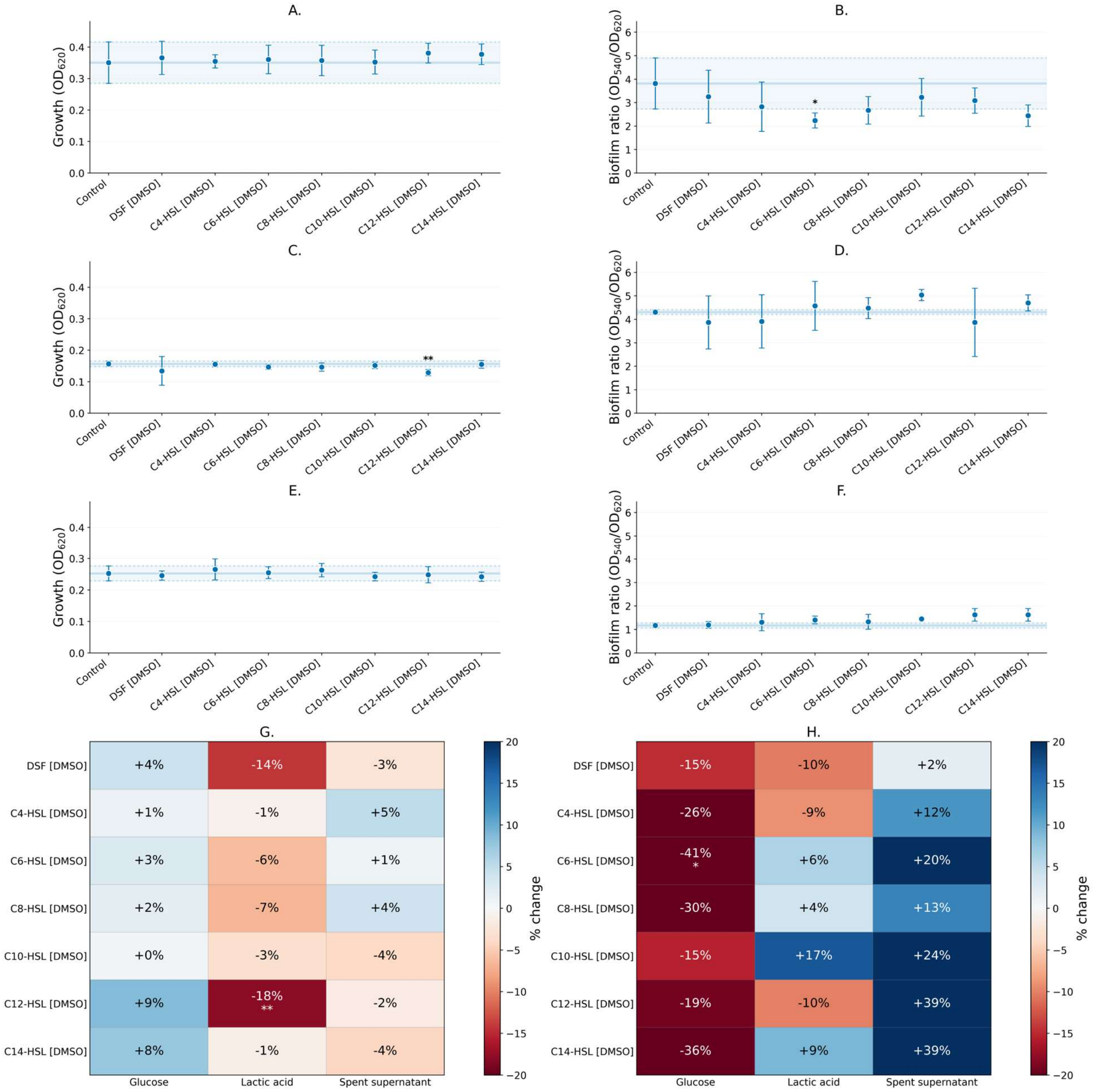

**Figure S 11. Growth (A, n=12) and biofilm ratio (B, n=8) of *L. plantarum* after 24 hours with different DMSO concentrations added corresponding to the quorum sensing conditions or without any DMSO added (control). Error bars indicate standard deviation. The solid line indicates the average value of the condition without exogenous quorum sensing molecules (control), while the dotted lines indicate the mean ± 1 standard deviation. Growth compared to the control (C) and biofilm ratio compared to the control (D) show the change in percent compared to the control. * indicates significant changes compared to the untreated control ($p < 0.05$).**

There was a limited effect on the growth, with some effect on the biofilm ratio. In three conditions (growth with the DMSO concentration of the condition with C12-HSL when growing on lactic acid, biofilm with the DMSO concentration of the condition with C6-HSL when grown on glucose and biofilm ratio with the DMSO concentration of the condition with the DMSO concentration of the condition with C10-HSL when grown on spent supernatant) are statistically significant. However, due to the low sample size (n=6 for growth, n=4 for biofilm ratio), the statistical interpretation is limited. Additionally, there is no clear DMSO dose-respondent response and the variance is especially for the biofilm quite large for some conditions. For this reason, DMSO was rejected as the explanation for the values observed in the experiment with exogenous quorum sensing molecules added.

#### S3.2 *M. elsdenii* experiments in serum bottles

The evolution of metabolites, the growth and pH can be seen in Figure S 12.

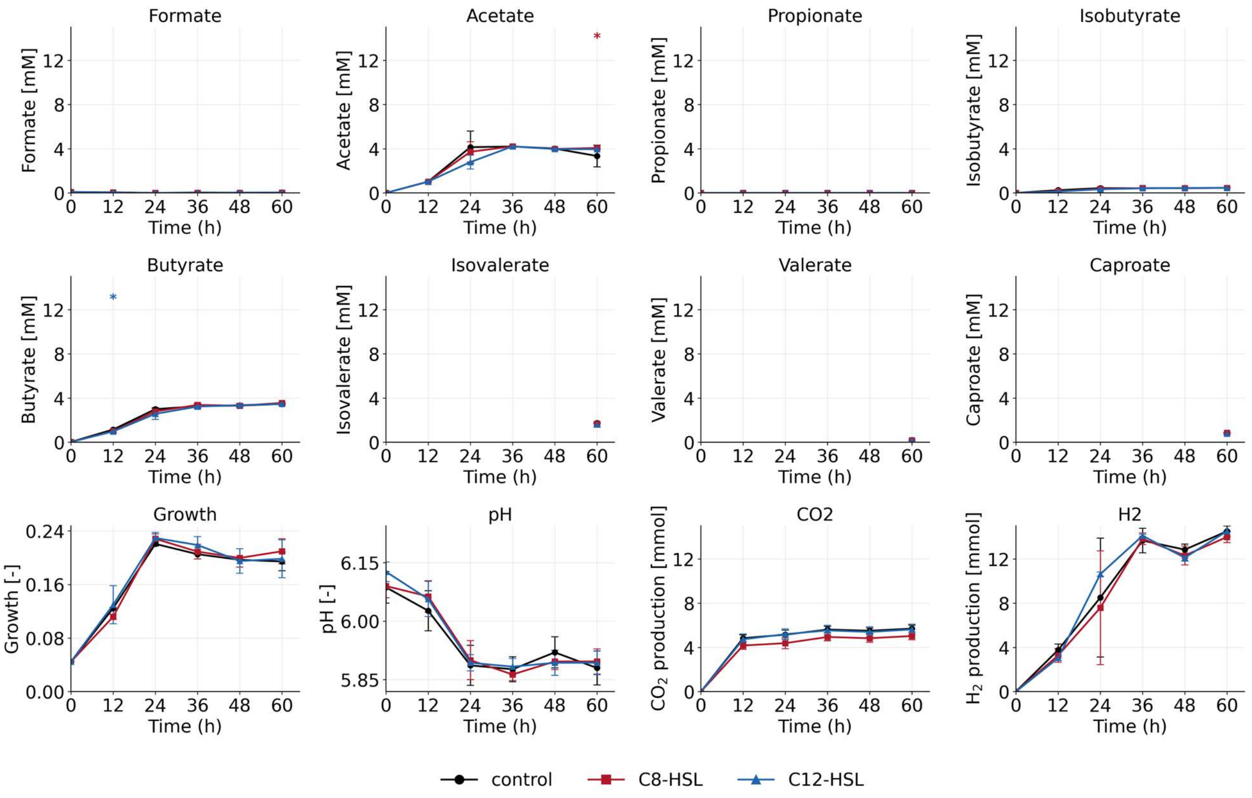

**Figure S 12. Concentration of acetate, propionate, isobutyrate, butyrate, isovalerate, valerate, caproate and growth in the serum bottles of M. elsdenii growing on glucose over time without added quorum sensing molecules (-•-), with added C8-HSL (-▪-) and with added C12-HSL (-▴-). * indicates significant values ($p < 0.05$)**

The electron balance of *M. elsdenii* growing on glucose for 60 hours in serum bottles can be seen in Figure S 13. All values included were based on measurements, not on calculations.

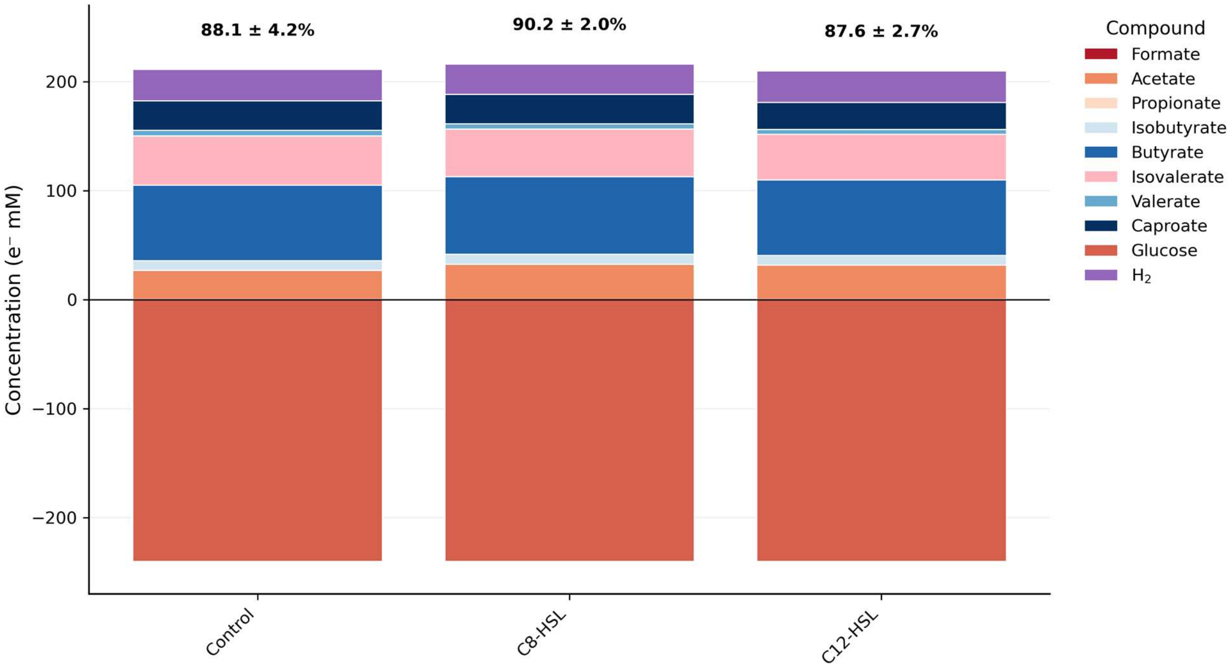

**Figure S 13. Electron balances of *M. elsdenii* grown for 60 hours on glucose without exogenous quorum sensing molecules (control) and with added quorum sensing molecules. Positive values indicate production, while negative values indicate consumption. Values above indicate the degree of closure (%) ± the standard deviation.**

### S4. Co-culture experiments

#### S4.1 Co-culture experiments in plates

The growth and biofilm ratio of the co-cultures in the plates at 24 hours and 48 hours can be seen in Table S 17 as the average value ± the standard deviation and the p-value in the brackets.

**Table S 17. Growth and normalized biofilm ratio of the *L. plantarum* and *M. elsdenii* co-culture after 24 hours and after 48 hours in 96 well plates. Statistically significant values ($p< 0.05$) using the steel-type test are indicated in bold (n=40) for growth and n=32 for biofilm)**

| | T = 24h | | T = 48h | |
|---|---|---|---|---|
| | Growth ($OD_{620}$) | Biofilm ratio ($OD_{540}/_{620}$) | Growth ($OD_{620}$) | Biofilm ratio ($OD_{540}/_{620}$) |
| control | 0.50 ± 0.03 | 0.80 ± 0.19 | 0.39 ± 0.04 | 3.11 ± 0.50 |
| +C8-HSL | **0.48 ± 0.03 (p = 0.04)** | 0.88 ± 0.25 (p = 0.22) | 0.39 ± 0.05 (p = 0.57) | 3.24 ± 0.52 (p = 0.31) |
| +C12-HSL | 0.49 ± 0.02 (p = 0.19) | **0.92 ± 0.18 (p = 0.04)** | **0.41 ± 0.03 (p = 0.01)** | 2.97 ± 0.41 (p = 0.46) |

The electron balances after 24 hours and 48 hours can be seen in Figure S 14A (24 hours) and Figure S 14B (48 hours).

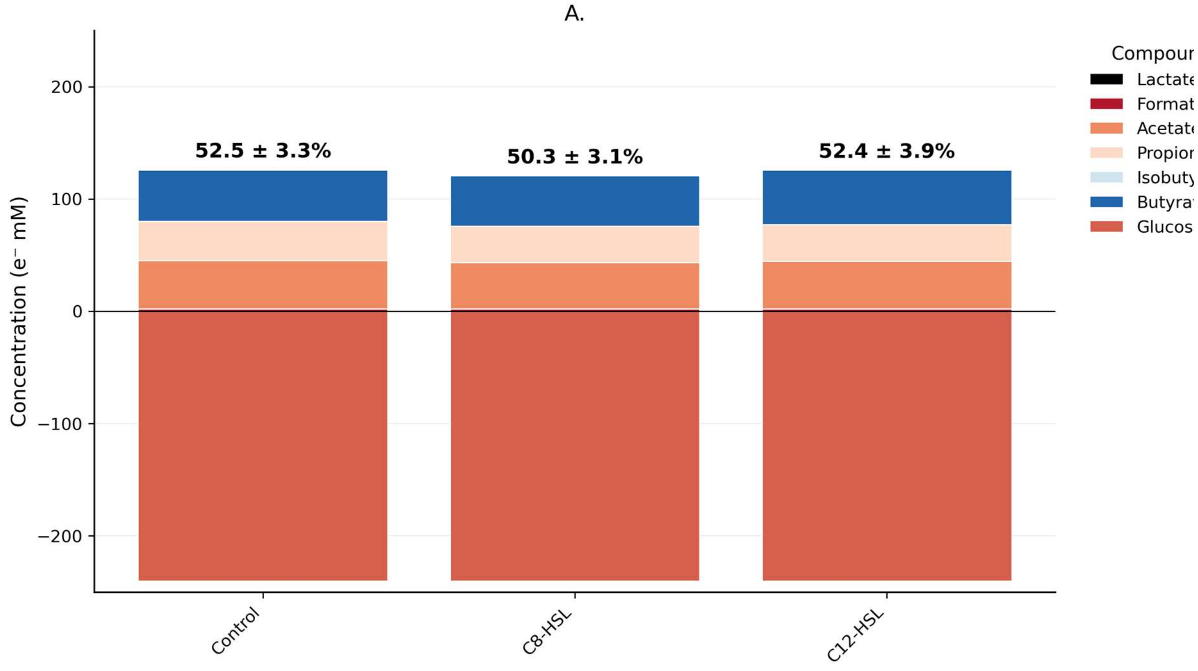

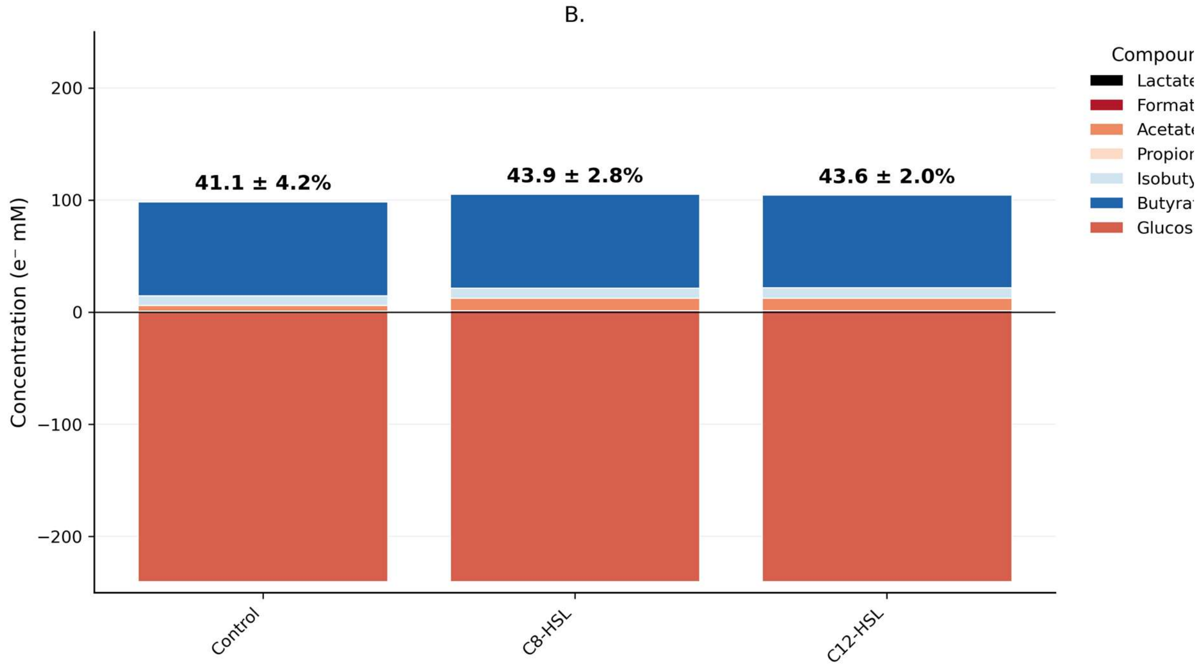

**Figure S 14. Electron balances of the co-culture of *L. plantarum* and *M. elsdenii* grown for 24 hours (A) and 48 hours (B) on glucose without exogenous quorum sensing molecules (control), with added C8-HSL and with added C12-HSL. Positive values indicate production, while negative values indicate consumption. Values above indicate the degree of closure (%) ± the standard deviation.**

At 24 hours, the electron balances closes for 52% for the control and the condition with exogenous C12-HSL and closes for 50% for the condition with exogenous C8-HSL. This reduces to 41%, 44% and 44% for the control, exogenous C8-HSL and exogenous C12-HSL respectively at 48 hours, indicating an increase in not-accounted for products. This is in line with growth on glucose or lactic acid, but not in line with the values obtained when growing on the spent supernatant.

#### S4.2 Co-culture experiments in serum bottles

The metabolites, growth and pH over time from the co-culture experiment in serum bottles can be found in Table S 18.

**Table S 18. Average values for the metabolite, growth ($OD_{620}$), pH and gas data with the standard deviation for each time point of the co-cultures growing in serum bottles. P-values were calculated using the steel-type max permutation test.**

| Time (h) | Condition | Glucose (mM) | Lactate (mM) | Formate (mM) | Acetate (mM) | Propionate (mM) | Iso-butyrate (mM) | Butyrate (mM) | Valerate (mM) | Iso-valerate (mM) | Caproate (mM) | Growth ($OD_{620}$) | pH (-) | H2 (mM) |
|---|---|---|---|---|---|---|---|---|---|---|---|---|---|---|
| 0 | Control | 8.61 ± 0.9 | 0.26 ± 0.01 | 0.08 ± 0 | 0 ± 0 | 0 ± 0 | 0 ± 0 | 0.03 ± 0 | 0 ± 0 | 0 ± 0 | 0 ± 0 | 0.04 ± 0 | 6.23 ± 0.02 | 0 ± 0 |
| 0 | C8-HSL | 6.94 ± 0.55 (p = 0.04) | 0.26 ± 0 (p = 0.99) | 0.08 ± 0 (p = 0.47) | 0 ± 0 (p = 1) | 0 ± 0 (p = 1) | 0 ± 0 (p = 1) | 0.03 ± 0 (p = 0.45) | 0 ± 0 (p = 0) | 0 ± 0 (p = 0) | 0 ± 0 (p = 1) | 0.04 ± 0 (p = 1) | 6.25 ± 0.03 (p = 0.66) | 0 ± 0 (p = 1) |
| 0 | C12-HSL | 7.16 ± 0.21 (p = 0.15) | 0.27 ± 0.01 (p = 0.4) | 0.08 ± 0 (p = 0.4) | 0 ± 0 (p = 1) | 0 ± 0 (p = 1) | 0 ± 0 (p = 1) | 0.03 ± 0 (p = 0.51) | 0 ± 0 (p = 0) | 0 ± 0 (p = 0) | 0 ± 0 (p = 1) | 0.04 ± 0 (p = 1) | 6.27 ± 0.06 (p = 0.56) | 0 ± 0 (p = 1) |
| 12 | Control | 5.15 ± 0.24 | 0.2 ± 0.09 | 0.1 ± 0 | 0.84 ± 0.01 | 0 ± 0 | 0 ± 0 | 0.79 ± 0.14 | 0 ± 0 | 0 ± 0 | 0.71 ± 0.71 | 0.08 ± 0 | 6.18 ± 0.03 | 1.72 ± 0.16 |
| 12 | C8-HSL | 5.15 ± 0.13 (p = 0.99) | 0.31 ± 0.14 (p = 0.51) | 0.1 ± 0 (p = 0.99) | 0.84 ± 0.05 (p = 0.81) | 0 ± 0 (p = 1) | 0 ± 0 (p = 1) | 0.71 ± 0.08 (p = 0.45) | 0 ± 0 (p = 0) | 0 ± 0 (p = 0) | 0.78 ± 0.78 (p = 0.27) | 0.08 ± 0 (p = 0.84) | 6.17 ± 0.02 (p = 0.51) | 1.6 ± 0.13 (p = 0.73) |
| 12 | C12-HSL | 5.13 ± 0.25 (p = 0.99) | 0.23 ± 0.1 (p = 0.75) | 0.1 ± 0 (p = 0.45) | 0.87 ± 0.03 (p = 0.46) | 0 ± 0 (p = 1) | 0.01 ± 0.01 (p = 1) | 0.76 ± 0.1 (p = 0.99) | 0 ± 0 (p = 0) | 0 ± 0 (p = 0) | 0.92 ± 0.92 (p = 0.75) | 0.08 ± 0 (p = 0.73) | 6.18 ± 0.01 (p = 1) | 1.3 ± 0.36 (p = 0.1) |
| 24 | Control | | 0.48 ± 0.46 | 0.14 ± 0.01 | 1.07 ± 0.06 | 0.04 ± 0.07 | 0.32 ± 0.02 | 3.25 ± 0.09 | 0 ± 0 (p = 0) | 0 ± 0 (p = 0) | | 0.19 ± 0.01 | 6.01 ± 0.03 | 4.56 ± 2.53 |
| 24 | C8-HSL | | 0.62 ± 0.15 (p = 0.87) | 0.11 ± 0.01 (p = 0.11) | 0.97 ± 0.1 (p = 0.57) | 0 ± 0 (p = 1) | 0.29 ± 0.05 (p = 0.65) | 3.03 ± 0.43 (p = 0.91) | 0 ± 0 (p = 0) | 0 ± 0 (p = 0) | | 0.2 ± 0.02 (p = 0.23) | 6.04 ± 0.01 (p = 0.36) | 5.35 ± 2.37 (p = 0.76) |
| 24 | C12-HSL | | 0.29 ± 0.31 (p = 0.71) | 0.12 ± 0.03 (p = 0.45) | 1.14 ± 0.39 (p = 1) | 0.25 ± 0.43 (p = 1) | 0.27 ± 0.09 (p = 0.65) | 2.98 ± 0.51 (p = 0.91) | 0 ± 0 (p = 0) | 0 ± 0 (p = 0) | | 0.2 ± 0.01 (p = 0.27) | 6.05 ± 0.01 (p = 0.04) | 5.62 ± 2.65 (p = 0.51) |
| 36 | Control | 0 ± 0 | 0 ± 0 | 0.19 ± 0.02 | 0.65 ± 0.05 | 0.08 ± 0.08 | 0.39 ± 0.05 | 3.43 ± 0.04 | 0 ± 0 (p = 0) | 0 ± 0 (p = 0) | 1.03 ± 1.03 ( | 0.2 ± 0.01 | 6.02 ± 0.03 | 9.09 ± 0.2 |
| 36 | C8-HSL | 0 ± 0 (p = 1) | 0 ± 0 (p = 1) | 0.17 ± 0.01 (p = 0.18) | 0.77 ± 0.2 (p = 0.4) | 0.27 ± 0.35 (p = 0.97) | 0.43 ± 0.04 (p = 0.26) | 3.49 ± 0.06 (p = 0.41) | 0 ± 0 (p = 0) | 0 ± 0 (p = 0) | 1 ± 1 (p = 0.91) | 0.21 ± 0 (p = 0.73) | 6.02 ± 0.03 (p = 0.98) | 9.18 ± 0.31 (p = 0.85) |
| 36 | C12-HSL | 0.33 ± 0.29 (p = 0.24) | 0 ± 0 (p = 1) | 0.18 ± 0 (p = 0.8) | 0.64 ± 0.01 (p = 0.99) | 0.02 ± 0.04 (p = 0.37) | 0.4 ± 0.01 (p = 0.93) | 3.45 ± 0.1 (p = 0.99) | 0 ± 0 (p = 0) | 0 ± 0 (p = 0) | 1.49 ± 1.49 (p = 0.14) | 0.2 ± 0 (p = 0.75) | 6.02 ± 0.01 (p = 1) | 9.42 ± 0.35 (p = 0.32) |
| 48 | Control | 0 ± 0 | 0 ± 0 | 0.21 ± 0.02 | 0.55 ± 0.06 | 0.04 ± 0.08 | 0.53 ± 0.02 | 3.74 ± 0.02 | 0 ± 0 | 0 ± 0 | 1.01 ± 1.01 | 0.17 ± 0.01 | 6.04 ± 0.06 | 7.74 ± 0.79 |
| 48 | C8-HSL | 0 ± 0 (p = 1) | 0 ± 0 (p = 1) | 0.18 ± 0.02 (p = 0.37) | 0.65 ± 0.18 (p = 0.31) | 0.2 ± 0.35 (p = 1) | 0.57 ± 0.1 (p = 0.97) | 3.77 ± 0.05 (p = 0.97) | 0 ± 0 (p = 0) | 0 ± 0 (p = 0) | 1.08 ± 1.08 (p = 0.32) | 0.17 ± 0 (p = 0.91) | 6.06 ± 0.04 (p = 0.81) | 7.91 ± 0.98 (p = 0.85) |
| 48 | C12-HSL | 0 ± 0 (p = 1) | 0 ± 0 (p = 1) | 0.2 ± 0.01 (p = 0.91) | 0.54 ± 0.01 (p = 0.84) | 0 ± 0 (p = 1) | 0.52 ± 0.01 (p = 0.99) | 3.78 ± 0.08 (p = 0.99) | 0 ± 0 (p = 0) | 0 ± 0 (p = 0) | 1.09 ± 1.09 (p = 0.15) | 0.17 ± 0.01 (p = 0.91) | 6.06 ± 0.02 (p = 0.92) | 8.66 ± 0.16 (p = 0.05) |
| 60 | Control | | 0 ± 0 | 0.23 ± 0.03 | 0.65 ± 0.24 | 0.1 ± 0.09 | 0.64 ± 0.01 | 3.97 ± 0.02 | 0.33 ± 0.07 | 2.39 ± 0.1 | 1.31 ± 1.31 | 0.16 ± 0.01 | 6.04 ± 0.03 | 10.29 ± 0.02 |

| | | | | | | | | | | | | | | |
|---|---|---|---|---|---|---|---|---|---|---|---|---|---|---|
| 60 | C8-HSL | | 0 ± 0 (p = 1) | 0.21 ± 0.02 (p = 0.29) | 0.57 ± 0.17 (p = 0.41) | 0.24 ± 0.34 (p = 1) | 0.68 ± 0.09 (p = 0.97) | 3.94 ± 0.12 (p = 0.74) | 0.49 ± 0.35 (p = 0.99) | 2.51 ± 0.15 (p = 0.73) | 1.27 ± 1.27 (p = 0.65) | 0.17 ± 0 (p = 0.47) | 6.06 ± 0.01 (p = 0.46) | 10.35 ± 0.42 (p = 0.65) |
| 60 | C12-HSL | | 0 ± 0 (p = 1) | 0.23 ± 0.01 (p = 0.98) | 0.47 ± 0.02 (p = 0.27) | 0.03 ± 0.06 (p = 0.55) | 0.63 ± 0.01 (p = 0.73) | 3.98 ± 0.09 (p = 0.85) | 0.24 ± 0.05 (p = 0.26) | 2.34 ± 0.09 (p = 0.75) | 1.33 ± 1.33 (p = 0.92) | 0.16 ± 0 (p = 0.99) | 6.05 ± 0.02 (p = 0.97) | 10.67 ± 0.31 (p = 0.15) |

The electron balance of the co-culture after 60 hours can be seen in Figure S 15. All shown compounds were measured, not calculated.

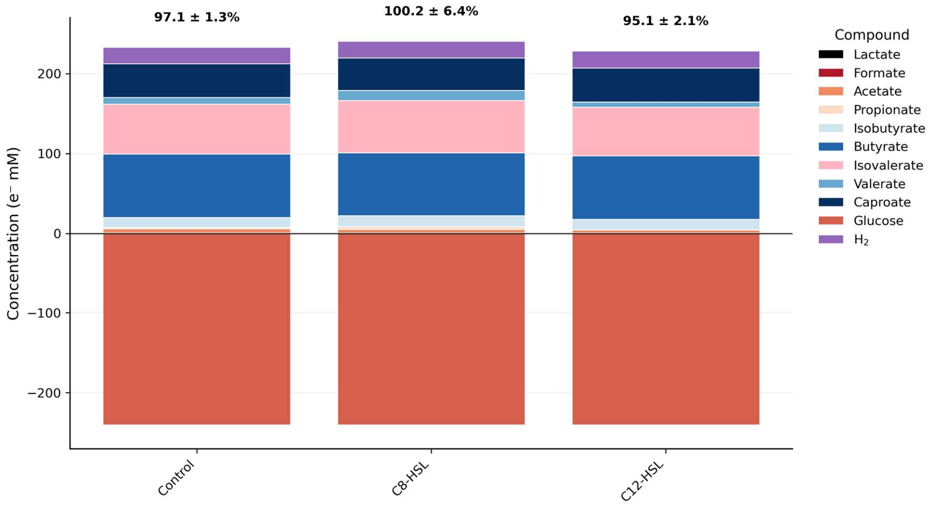

**Figure S 15. Electron balance of the co-culture after 60 hours growing in serum bottles on glucose.**

The values for the soluble EPS, bound EPS and biomass can be found in Table S 19.

**Table S 19. Soluble EPS, bound EPS and biomass measured after 60 hours. Values that showed an increase compared to the control are indicated in blue, while values that showed a decrease compared to the control are indicated in red. Statistically significant values ($p < 0.05$) are indicated in bold (n=3).**

| | Soluble EPS ($mg_{COD}$/L) | Bound EPS ($mg_{COD}$/L) | Biomass ($mg_{COD}$/L) |
|---|---|---|---|
| *L. plantarum* | 516.45 ± 228.40 | 213.33 ± 12.04 | 109.07 ± 21.19 |
| *M. elsdenii* on glucose | 440.66 ± 345.80 | 332.67 ± 99.28 | 198.60 ± 16.85 |
| Co-culture | 276.18 ± 23.03 | 291.33 ± 29.04 | 113.80 ± 10.31 |
| Co-culture + C8-HSL | 352.95 ± 577.56 | 264.00 ± 23.04 | 101.13 ± 3.37 |
| Co-culture + C12-HSL | 610.21 ± 214.26 | 304.67 ± 32.22 | 75.00 ± 26.73 |

The quorum sensing inhibition based on the detector strain *Chromobacterium violaecum* ATCC 12472 can be found in Table S 20. *C. violaceum* ATCC 12472 naturally produces and is induced by long-chain AHLs (including C12-HSL) and is inhibited by short-chain AHLs (Morohoshi et al. 2008). The negative value for the conditions with added C12-HSL and the very high positive value for the conditions with added C8-HSL therefore are a rough indication of the presence of C12-HSL and C8-HSL in the media. For *L. plantarum*, the quorum sensing inhibition is 17.19%, while for *M. elsdenii* the quorum sensing inhibition is 9.06%. However, for the co-culture, the quorum sensing inhibition is only 2%, indicating that there is less inhibition in the co-culture compared to either of the pure culture.

**Table S 20. QSI determination of the pure cultures and the co-cultures with or without exogenous quorum sensing molecule[illegible] (n=3 biological, n=3 technical)**

| | Violaceum production ($OD_{584}$) | QSI |
|---|---|---|
| *L. plantarum* | 1.951 ± 0.125 | 17.19 |
| *L. plantarum* + C8-HSL | 1.331 ± 0.211 | 43.51 |
| *M. elsdenii* | 2.143 ± 0.081 | 9.06 |
| *M. elsdenii* + C8-HSL | 1.337 ± 0.097 | 43.26 |
| *M. elsdenii* + C12-HSL | 2.432 ± 0.525 | -3.23 |
| *L. plantarum* and *M. elsdenii* co-culture | 2.309 ± 0.344 | 2.01 |
| *L. plantarum* and *M. elsdenii* co-culture + C8-HSL | 1.465 ± 0.081 | 37.82 |
| *L. plantarum* and *M. elsdenii* co-culture + C12-HSL | 2.414 ± 0.228 | -2.47 |
| Control (media only) | 2.356 ± 0.126 | - |
